# Long-read sequencing reveals genomic landscape of structural variants in a major disease vector

**DOI:** 10.64898/2026.10.05.756236

**Authors:** Remi N. Ketchum, Anna W. Trotter, Rafael J. Vivero, Madelyn A. Pivovarnik, Daniel R. Schrider, Daniel R. Matute

## Abstract

The southern house mosquito *Culex quinquefasciatus* is a major disease vector for multiple arboviruses, including West Nile virus and St. Louis encephalitis virus. Insecticides are widely used to control *Cx. quinquefasciatus* populations, but resistant mosquitoes have emerged in numerous locations across the globe, including the United States. Much progress has been made in recent years in identifying insecticide resistance (IR) mutations in various pest species. In most species, these studies have focused primarily on SNPs as they are readily detectable. However, several genomic structural variants (SVs) have been experimentally shown to impact IR in *Drosophila melanogaster*, and structural variation has been observed in natural populations of vector species at a few genes that may be relevant to IR (e.g., cytochrome P450s). Because most population genomic data are derived from short sequencing reads, which fail to detect a large fraction of SVs, the contribution of structural variants to the evolution of IR and other adaptations in vector species is likely to be underestimated in the absence of long-read sequencing data. Here, we present the first genome-wide assessment of structural variants in *Cx. quinquefasciatus* from wild populations in Puerto Rico and Zambia using PacBio long-read sequencing. We constructed high-quality genome assemblies for each and found a total of 259,046 SVs which consisted predominantly of insertions and deletions. We found that 154,071 of these variants contained or overlapped genes. Of these SVs, 5,225 intersected IR genes and 56 of these were predicted to result in changes to downstream protein function. Several SVs predicted to severely impact protein-coding genes also coincided with signatures of selective sweeps, highlighting their possible role as targets of selection. Many of these variants would have gone undetected using short-read approaches, underscoring the importance of long-read sequencing for comprehensive population genetic analyses of adaptation to pesticides and other strong selective pressures, not only in insects but across the tree of life.

## Introduction

*Culex quinquefasciatus,* or the southern house mosquito, is a widely distributed tropical and sub-tropical mosquito that is part of the *Cx. pipiens* complex [1]. Although it remains unclear whether *Cx. quinquefasciatus* originated in East Africa or Southeast Asia [2–4], it has subsequently radiated outwards, aided by historical and contemporary human migration [5]. The contemporary distribution of *Cx. quinquefasciatus* includes Australia, Polynesia, North and South America, South and East Asia, and Africa [3,4] and is continuing to expand as climate change increases temperatures at higher latitudes [6,7]. The species is responsible for the transmission of several human arboviruses that cause diseases such as the St. Louis encephalitis virus, West Nile virus, and Rift Valley fever virus [8]. The World Health Organization (WHO) estimated that there were 100,000 clinical cases resulting from pathogens transmitted by *Culex* spp. and global modeling estimated a total of 25,000 deaths in 2015 [9]. *Culex quinquefasciatus* breeds preferably in stagnant, polluted water sources found in urban areas, including plastic containers, tires, and drains, which leads to increased disease transmission in heavily populated areas [10–12]. The species’ vectorial capacity and ability to thrive in urban environments make *Cx. quinquefasciatus* a significant threat to human health [7]. Although *Cx. quinquefasciatus* is a major disease vector and is known to adapt rapidly to changing environmental conditions and human interventions such as insecticide use [13–15], we know little about the genomic features that underlie this adaptation.

To curb population growth of *Cx. quinquefasciatus* and other mosquito species, and thereby slow the spread of disease, governments have employed insecticide treatment campaigns in several regions around the world [8,16,17]. Historically, pyrethroids and organophosphates are among the most common classes of insecticides used in these campaigns [18]. Pyrethroids act by targeting voltage-gated sodium channels, thereby disrupting nerve impulse transmission and eventually leading to paralysis and death [19,20]. Similarly, organophosphates inhibit the acetylcholinesterase enzyme, which leads to an abundance of acetylcholine and subsequent disruption of neurotransmission [21,22]. Although these treatments are effective initially when applied to non-resistant populations, many insects—including *Cx. quinquefasciatus*—quickly evolve resistance, eventually rendering the insecticides significantly less effective. Indeed, *Culex* mosquitoes tend to exhibit higher levels of insecticide resistance than more well-studied species of mosquitoes like *Aedes aegypti* in regions where they coexist [15]. Resistance to both pyrethroids and organophosphates has been reported in *Cx. quinquefasciatus* from the United States [15,23], Brazil [8], and Thailand [24] and resistance to pyrethroids has been reported in Benin [25] and West Africa [26]. This resistance typically manifests through behavioral changes, variation in genes encoding for insecticide target-sites (e.g., the *voltage gated sodium channel* for pyrethroids or *acetylcholinesterase*-*1* genes for organophosphates), changes to gene expression levels associated with detoxification (e.g., cytochrome P450s or glutathione *S-*transferase genes), or with cuticular thickening or changes in composition [27–30].

Insecticide resistance provides an excellent model for studying rapid adaptation because insects often have the biological traits which promote fast responses to new selective pressures: large population sizes [31–33], short generation times [34,35], and high reproductive rates. Further, insecticides can eliminate a substantial proportion of a population, thereby creating the strong selective pressure that is required to drive rapid evolutionary change. However, research in mosquitoes has largely focused on investigating adaptive signatures associated with single nucleotide polymorphisms (SNPs) [36–41] and has consequently overlooked larger structural variants (SVs) as they are much more challenging to genotype using short-read sequencing (but see: [42–45]). Genomic SVs, or genome modifications that are larger than 50 bp and include inversions, insertions, duplications and deletions, often account for a greater proportion of genomic sequence variation than SNPs [46–48]. They also tend to have larger fitness effects because they are more likely to disrupt—or even contain—entire genes and other functional elements [49]. Such large-effect mutations may often help facilitate adaptation in populations that are subject to a severe change in the selective environment such that they are now located far from their phenotypic optimum [50,51]. As insecticide resistance poses a large public health threat and is one of the most important impediments to the chemical control of insect pests, understanding the role of SVs in the evolution of IR is crucial.

Structural variants have been implicated as a major contributor to rapid adaptation in a wide range of species [52]. For example, in *Drosophila* species there are latitudinal chromosomal inversions that are responsible for seasonal adaptation and differences in body size [53–55]. In the peppered moth, a large transposable element insertion resulted in the replacement of the common pale typica form by the black form in Britain during the Industrial Revolution [56]. In snakes [57], spiders [58], gastropods [59], and sea anemones [60], gene duplications have been implicated in venom complexity and expression. While loss-of-function mutations such as deletions were historically thought to be largely deleterious (outside of the removal of gene redundancy), recent studies have revealed that gene loss can be an important driver of adaptation [61–66]. In *Anopheles* mosquitoes, deletions of redundant gene copies reduced the fitness costs of gene duplication while maintaining the essential insecticide resistance gene, *ace-1* [67]. Further, deletions were shown to improve the speed and capacity for adaptation to future stressors in yeast [62]. These examples (as well as many more, see review [65]) support the idea that gene loss can act as an adaptive evolutionary mechanism.

The proliferation of large genomic rearrangements can be a result of regions rich in TEs or repeats which can confound cellular machinery due to their high sequence similarity [68]. This can happen through replication slippage, where DNA polymerase pauses on repetitive sequences and the newly made strand separates and misaligns, ultimately resulting in either insertions or deletions [69]. Alternatively, non-allelic homologous recombination (NAHR), where chromosomes misalign during cell division, can also lead to the development of SVs [68]. Most studies investigating SVs and adaptation have focused on model species such as *Drosophila* [70,71], which do not have TE- or repeat-rich genomes when compared to mosquitoes from the subfamily Culiciniae (recent estimates of TE content range from 52.7–54.4% across the *Cx. pipiens* complex [72]). The repeat content in *Culex quinquefasciatus*, for example, comprises about 58% of the genome compared to only about 22% in *D. melanogaster* [73,74]. Although high levels of repeat content can facilitate SV formation, they can also hinder SV detection due to the unreliable mapping of short reads [75]. Short-read data also often suffers from high false-positive and false-negative rates and inaccurate breakpoint resolutions of genomic regions [76,77]. This results in a situation where organisms that are the most challenging to study are also those where an examination of SVs is most warranted. Thankfully, advances in long-read sequencing provide unprecedented opportunities to study SVs as such reads can span SVs and their breakpoints and facilitate more contiguous assemblies [77–79].

Given that *Culex* mosquito genomes contain genomic features that promote structural variation, and that these species continue to expand their ranges and demonstrate remarkable capacity for rapid adaptation and disease transmission, they provide an important model for understanding the evolutionary impact of SVs. However, only a handful of studies have detected SVs in IR genes (namely *vgsc* [80], cytochrome P450s [81–84], and *ace-1* [85]) in *Cx. quinquefasciatus*. Although long-read sequencing has the capacity to identify previously intractable genomic rearrangements, no studies to date have used long-reads to investigate SVs in *Cx. quinquefasciatus* (however, see [72] for a between-species investigation of copy number variation). This represents a significant knowledge gap, given the species’ potential to become a considerable public health burden on a global scale. Here, we use long-read sequencing to uncover SVs in *Cx. quinquefasciatus* population samples from Zambia and Puerto Rico for the first time. We describe the diversity and distribution of SVs in these population samples and highlight key SVs that are likely to have large functional impacts on protein-coding genes. As these populations are under strong insecticide pressures [86–88], we also highlight SVs that may be involved in adaptation to insecticides, as well as other environmental changes.

## Materials and Methods

### Sampling, DNA Extraction, and Sequencing

In 2019, we collected four *Cx. quinquefasciatus* mosquitoes from Kakumbi by the Luangwa River (Zambia) and three mosquitoes in Puerto Rico using a mechanical aspirator from mosquitoes resting on walls. The sorting and identification procedure was identical for the two locations. Mosquitoes were sorted with a Leica microscope in the field and assigned to morphospecies and sex. Specimens were kept in 90% molecular biology grade ethanol and imported into the United States. To extract DNA, we used mechanical lysis of individual specimens by adding 0.5 mm diameter zirconia/silica beads and lysis buffer to the preheated tissue and vigorously agitated the mixture using a Qiagen Vortex for 15 minutes. To purify DNA, we used the DNeasy Blood and Tissue Kit (Qiagen, Hilden, Germany) according to the manufacturer’s standard protocol. Finally, we evaluated DNA integrity and concentration using agarose-gel electrophoresis (0.8% w/v) and Qubit fluorimetry (Qubit fluorimeter, Thermo Fisher Scientific, Waltham, MA), respectively. Extracted high-molecular-weight gDNA was shipped to the Genomics Core Facility at the Icahn School of Medicine at Mount Sinai, where libraries were prepared using PacBio’s Ultralow Input HiFi sequencing protocol (Pacific Biosciences, Menlo Park, CA, USA).

### Genome Assembly and Annotation

In order to generate seven high-quality genome assemblies, we first constructed initial assemblies using hifiasm v0.25.0 [89] with default parameters. To reduce the percentage of duplication present in our assemblies, we ran PurgeHaplotigs v1.1.3 [90] and set our percent cutoff for identifying a contig as a haplotig to 50%, as this yielded the most complete assemblies. These assemblies were then run through the PurgeHaplotigs ‘clip’ option to identify and trim overlapping contig ends. The K-mer Analysis Toolkit (KAT) v2.4.1 [91] was used to assess whether or not our assemblies were sufficiently purged of false duplications. These plots revealed that our assemblies were not sufficiently collapsed so we then purged our assemblies using purge_dups v1.2.6 [92] with default parameters. Finally, ragtag v2.1.0 [93] was used to correct and scaffold each assembly based on the *Cx. quinquefasciatus* reference genome (NCBI accession GCF_015732765.1), and seqkit v2.10.0 [94] was used to remove scaffolds shorter than 1,000 bp. Genome assembly and annotation qualities were assessed with BUSCO v6.0.0 [95] using the diptera_odb12 lineage dataset [95].

### Species Confirmation

Species within the *Culex* genus can be difficult to distinguish through visual assessment alone [96]. We therefore corroborated our morphological identification with both mitochondrial and nuclear phylogenies. First, we assembled mitochondrial genomes for each sample using MitoHifi v3.2.2 [97], using the raw HiFi reads as input. Second, we downloaded 29 *Culex* and one *Aedes aegypti* mitochondrial genomes from NCBI to include in our analysis. Each genome was rotated to the same start position using rotate v1.0 (https://github.com/richarddurbin/rotate) with the anchor seed set to “TCGCGACAATG.” This anchor was chosen as it is part of the *COX1* gene which is highly conserved across *Culex* species [98]. Third, we aligned our 37 sequences with a MAFFT alignment and inferred the phylogenetic tree using UGENE [99]. IQ-TREE v2.4.0 [100] was used to infer a maximum likelihood tree from this rotated alignment. The tree was generated with the GTR+F+I+R3 model selected by ModelFinder [101], and branch support was evaluated with 1,000 ultrafast bootstraps using the option “-B 1000” [102]. Finally, iTOL v7 [103] was used to visualize the tree with *Ae. aegypti* set as the outgroup.

To generate a species tree for the nuclear genome, we employed the deep learning algorithm Helixer v.0.3.4 [104,105] for *de novo* genome annotation of our genome assemblies and several *Culex* genome assemblies available through NCBI. These included: *Cx. modestus* (GCA_964213205.1), *Cx. perexiguus* (GCA_964243045.1), *Cx. theileri* (GCA_964340705.1), *Cx. laticinctus* (GCA_964187835.1), *Cx. pipiens pipiens* (GCA_963924435.1), *Cx. pipiens pallens* (GCF_016801865.2), *Cx. quinquefasciatus* (GCF_015732765.1), and *Cx. tarsalis* [106–110]. Gffread v.2.4.0 [111] was used to convert the Helixer GFF output into a protein sequence file for each individual. For *Cx. quinquefasciatus* and *Cx. pipiens pallens*, protein sequences were downloaded directly from NCBI. The protein FASTA files were used for orthology inference and species tree reconstruction using the multiple sequence alignment mode in OrthoFinder v3.1.5 [112]. The resulting SpeciesTreeAlignment.fa file was input into IQ-TREE v2.4.0 [100], which inferred a maximum likelihood phylogenetic tree from the concatenated protein alignment using the JTT+F+R10 model selected by ModelFinder [101]. Branch support was evaluated with 1,000 ultrafast bootstrap replicates using the option “-B 1000” [102]. The resulting tree was rooted at the outgroup *Cx. tarsalis* and visualized using iTOL.

### Structural Variant Calling

To generate a high-quality consensus set of structural variants, we used a combination of both read- and assembly-based callers. For the read-based callers, we first aligned our raw reads to the published *Cx. quinquefasciatus* genome using pbmm2 within minimap2 v.2.30 to generate BAM files for each sample [113]. Our read-based callers included cuteSV v2.1.3 [114], Sniffles2 v.2.0.7 [115], and pbsv v2.11.0 (https://github.com/PacificBiosciences/pbsv), while our assembly-based caller was SVIMasm v1.0.3 [116]. CuteSV was run with the recommended parameters for PacBio HiFi reads. This includes setting the maximum distance for clustering reads into the same insertion or deletion event to 1,000 bp. Additionally, we set the base pair identity ratio to 0.9 for insertions and 0.5 for deletions. This means that if the fraction of matching base pairs between sequencing reads and alignment breakpoints is above these values, the breakpoints are not merged. For insertions, this means that insertions with a length difference of less than the average insertion length within a region are considered to represent the same insertion event. For deletions, the difference in length between events must be less than 0.5x the average length to be merged. We also set the maximum number of split segments a read may be aligned to before it is ignored to four instead of the default seven to curate a more stringent SV dataset. We next ran Sniffles with the minimum number of supporting reads needed for an SV to be reported set to eight and the minimum length needed for a deletion to be subject to central coverage drop-based filtering to 10,000 bp. Specifically, the mean coverage within the 500 bp before and after the deletion was calculated. If the mean coverage within the deletion was greater than 2/3 of the mean coverage of the surrounding regions, the deletion was removed. We also performed split-read filtering by setting the number of splits per kilobase of read sequence allowed to 0.1 and the base number of splits allowed to two. pbsv was run using default parameters for HiFi sequencing. Finally, we ran SVIM-asm by first mapping our assembled genomes to the *Cx. quinquefasciatus* reference genome using minimap2 with options “-ax asm5 -r2k,” to indicate assembly-to-assembly mapping and use a bandwidth of 2,000 bp to improve long indel alignments. The resulting BAM files were sorted and indexed with samtools v1.23.1 [117]. Then, SVIM-asm was run with default parameters for haploid genomes. Each of these four tools was run on each sample separately.

After SVs were called using these tools, the results for each sample individually were combined with combiSV v2.3, using default parameters [118]. The combiSV script was modified to remove logic that required a minimum number of supporting reads for SVIM SVs, as the assembly-based caller SVIM-asm was used in place of the traditional read-based SVIM. Each VCF file was filtered to retain variants that were supported by at least two SV callers. We then used Jasmine v1.1.5 [119] to merge all individual sample VCFs into a single combined VCF. Subsequently, BCFtools v1.23.1 was used to split the merged VCF back into per-sample VCFs, enabling a genotyping step using Sniffles. After genotyping, the resulting variant calls for each sample were merged back into a single VCF, producing a final dataset with genotypes assigned to all structural variants. This VCF was then filtered to remove SVs that were homozygous for the reference in every individual and SVs with length >100,000 bp. These larger SVs were removed after an analysis of the SV length distribution and manual examination of read depth found a high proportion of apparent false positives above this length cutoff (Supplemental Figure 1). This final dataset, which we will herein refer to as the full set of SVs, was used for all downstream analyses.

### SNP Calling

SNPs were jointly called across all seven *Cx. quinquefasciatus* samples using BCFtools v1.24 with the PacBio CCS configuration flag. To do so, we aligned the reads to the *Cx. quinquefasciatus* reference genome, variants were called using BCFtools mpileup and call, and we set our maximum read depth to 1,000. Variants were filtered to only retain sites which had quality scores greater than or equal to 20, and we restricted our filtering to only include SNPs (as the BCFtools calling method includes INDELs). For PCA, the SNP dataset was further filtered to only include biallelic SNPs that had been genotyped in every individual and had a minor allele count of at least three. Allele frequencies were calculated in PLINK v2 [120] and used as input to generate PCA plots.

### Functional Annotation of Variants

SnpEff v5.4c [121] was used to annotate SVs and predict their potential effects on genes and transcripts. Variants were classified according to the predicted severity of their effects, based on their genomic location and overlap with annotated genomic features from the published *Cx. quinquefasciatus* genome. The gene annotation used for SnpEff was downloaded as a GTF file from NCBI accession GCF_015732765.1 on March 6, 2026. To generate a dataset exclusively containing SVs predicted to have a significant downstream functional impact, we filtered the full set of SVs for all SVs labelled with the “high” functional impact annotation. This dataset is referred to as the high impact set of SVs. The high functional category included variants labeled as: chromosome number variation, rare amino acid variant (neither of which were present in our dataset), exon loss variant, frameshift variant, splice acceptor variant, splice donor variant, start lost, stop gained, stop lost, and transcript ablation. Importantly, each SV can have multiple SnpEff annotations with different predicted impacts. If an SV receives any annotations in the high category, it is categorized as high, otherwise if it receives any annotations in the “modifier” or “moderate-impact” or “low-impact” category, it is assigned to that respective category. We performed the same analysis using SNPs. For each genome, we also investigated the number of loss-of-function SNPs and SVs identified by SnpEff. We defined LOF variants as any variant containing the annotation “LOF” for all transcripts of at least one gene. We calculated the number of LOF variants with at least one non-reference allele, and the number of homozygous variants for each sample.

### Population Structure and Allele Frequency Calculations

To assess population structure based on the high-quality full set VCF, we used principal component analysis (PCA). Variants with missing genotype calls were excluded from the dataset using BCFtools v1.24 by filtering out sites that contained missing genotypes (GT= “mis”). We calculated allele frequencies for all variants and computed the first seven principal components using PLINK v2.0. The resulting PCA was plotted in ggplot2 in RStudio [122]. The VCF excluding missing genotypes was also used to generate the site frequency spectra of SVs for the two populations.

We then examined the extent and direction of allele frequency divergence between the two populations. For each population, alternate allele frequency was calculated as the proportion of alternate alleles among all non-missing alleles. We assumed diploid genotypes, with homozygous reference, heterozygous, and homozygous alternate genotypes contributing 0, 1, and 2 alternate alleles, respectively. Based on this distribution, we calculated the 5th (PR-enriched) and 95th (LZV-enriched) percentile thresholds and extracted each SV below and above those thresholds, respectively (Supplemental Figure 2). We will hereafter refer to these datasets as the PR-enriched and LZV-enriched set of SVs. Finally, we also generated a dataset which included both the SVs present in the PR-enriched VCF and LZV-enriched VCF, which we will refer to as the LZV/PR-enriched set of SVs.

### Genomic Regions under Positive Selection

We used SweepFinder2 [123] to identify regions of the *Cx. quinquefasciatus* genome under positive selection. We used publicly available Illumina sequencing data for *Cx. quinquefasciatus* samples from [124](NCBI BioProject accession: PRJNA1209100). All samples with mean genome-wide sequencing coverage less than 10x were removed. Alignment and variant calling were carried out following the workflow described in https://github.com/YukiHaba/PipPop_molestus_origin/tree/main. Briefly, we aligned read data to the *Cx. quinquefasciatus* reference genome using BWA-MEM2 [125], then marked duplicates using Picard (https://broadinstitute.github.io/picard/) and performed local realignment around indels with GATK [126]. SNPs were called using BCFtools. We also followed their SNP filtering criteria, which required that SNPs were biallelic, had a variant quality score >30, a mapping quality score >40, had <50% missingness across all samples (modified from 25% in the original study), and were within accessible, non-repetitive genomic regions. Specifically, repeat elements identified by RepeatMasker as described in [127] and sites with <0.5X or >1.5X normalized coverage were masked (see detailed methods described in Haba et al. 2025. Samples from three Sub-Saharan African locations were considered: South Africa (n=16), Madagascar (n=9), and Cameroon and Gabon (n=9). Cameroon and Gabon were considered as one population sample based on the results of the PCA (Supplemental Figure 3). These population samples were chosen because they were geographically closest to our Zambian population sample, the rationale being that some of the selective pressures faced by these samples may be more similar to those faced by the Zambian sample. The resulting VCF files were used to generate the input to SweepFinder2 with a custom python script (https://github.com/SchriderLab/timesweeper-experiments/blob/c674c9ebf44399af24b2475af994d4d640f08cb9/scripts/comp_methods/vcf2sf.py). The resulting allele frequency files were used to calculate the empirical site frequency spectrum which was then input as the background SFS in SweepFinder2. SweepFinder2 was run for each chromosome with a grid spacing of 1,000 bp and selective sweeps were defined as stretches of consecutive sites in the grid where SweepFinder identified a composite likelihood ratio (CLR) above the 99^th^ percentile for that chromosome. Note that we were unable to perform a similar analysis for samples located near Puerto Rico as Haba et al.’s (2025) sampling sites were all geographically very distant from this location. Using the selective sweep regions inferred here, we generated a dataset which contained SVs that overlapped selective sweeps, which we refer to as the sweep set of SVs. Finally, we obtained the intersection between the high impact set of SVs and the sweep set, and we refer to this as the high impact sweep set of SVs.

*Insecticide Resistance Gene List Curation.* To identify candidate insecticide resistance genes in *Cx. quinquefasciatus*, we used OrthoFinder to identify orthologs of the *Anopheles gambiae* IR genes curated by the *An. gambiae* 1000 Genomes Project [128]. The proteomes used as input for OrthoFinder were obtained from Ensembl Metazoa for *Ae. aegypti* (GCA_002204515.1) and *An. gambiae* (GCA_000005575.1), and from NCBI for *Cx. quinquefasciatus* (GCF_015732765.1)[127,129–131]. Using the OrthoFinder output, we identified orthogroups containing transcripts corresponding to *An. gambiae* IR gene products. Candidate IR genes in *Cx. quinquefasciatus* were defined as those with transcripts within these orthogroups, representing putative orthologs of *An. gambiae* IR-associated proteins. Corresponding gene IDs were then retrieved using NCBI’s datasets tool [132](Supplemental Table 1). To generate VCF files for analyzing SV impacts on IR genes, we filtered each of the seven final SV VCF files for SVs overlapping putative IR genes.

### Environmental Gene List Curation

In addition to IR genes, we investigated the impact of SVs on genes that are implicated in environmental sensing and adaptation in mosquitoes (Supplemental Table 2). We followed the method used above for IR genes to identify *Cx. Quinquefasciatus* orthologs of the *Ae. aegypti* chemosensory genes curated in [133]. We examined the list of genes overlapping high impact SVs for these orthologs, as well as any odorant receptor, heat shock, ionotropic receptor, and immune genes, as these gene families are known to mediate environmental interactions in mosquitoes [133–139].

### Enrichment Analysis

To look for overrepresented GO terms among genes overlapping structural variants in the seven different datasets (this included: the full set, high impact, sweep, high impact sweep, the PR-and LZV-enriched, and the LZV/PR-enriched sets of SVs) we performed a permutation-based enrichment test. We generated an input file from the full set of SVs and then treated each of the three chromosomes as a circular sequence. For each of the 10,000 permutations, a random circular offset was independently selected for each chromosome and then applied to all SV coordinates on that chromosome. In this way, we preserved the relative spacing between the SVs while simultaneously changing their genomic position. SVs that cross the chromosome boundary following rotation were split into two intervals but retained their original SV identifier. Next, we compared the number of genes belonging to each GO category that were within or overlapping the observed SVs against the corresponding number within the permuted SV set. A GO term was considered enriched when the observed SV set overlapped significantly more genes from that category than in the permuted set. To account for the proportion of true null hypotheses among the tested GO terms, we applied an adaptive false discovery rate correction based on Storey’s method [140]. Empirical *P*-values were calculated for each GO term as the proportion of permuted datasets in which the number of genes overlapping the term was greater than or equal to the number observed in the real data. The proportion of true nulls (π₀) was then estimated using a fixed tuning parameter of λ = 0.5, as the number of GO terms with P > 0.5 divided by m(1 − λ), where m is the number of tested terms, with π₀ capped at 1. Q-values were obtained by scaling *P*-values by m and π₀, dividing by the number of terms with an equal or smaller *P*-value, and enforcing monotonicity across terms. For terms with no permuted dataset equalling or exceeding the real overlap, the q-value was reported as an upper bound calculated from a minimum *P*-value of 1/N, where N is the number of permutations. Because a large number of GO terms had few or no genes overlapping the real significant gene list, many of the permuted datasets matched or exceeded this minimal real overlap, giving these terms *P*-values of 1. We note that multiple testing corrections were applied separately within each dataset but not across all datasets meaning that the overall number of false discoveries across the entire collection of datasets may remain elevated.

We then used this permutation test to assess whether our SV regions were enriched for overlap with genes from the curated IR gene set described above. For the observed data, each IR gene was classified as overlapping an SV region if any part of the gene interval intersected an SV. We extended the gene windows by 1 kb in either direction to account for nearby regulatory elements. Statistical significance was assessed using the circular-shift permutations described above where the number of IR genes overlapping each of the 10,000 permutations was calculated, forming our null distribution of overlap counts. Finally, we investigated whether SV regions were enriched for selective sweeps by permuting SV regions for the LZV-enriched VCF (as the sweep regions identified here reflect selection detected in sub-Saharan Africa) and repeating the above analysis using a concatenation of the CLR outlier regions for each population and chromosome.

### Global prevalence of GST deletion

To explore the global prevalence of *GST-1* deletions we discovered within our data (as described in the Results), we used alignments generated for our SweepFinder analysis from the *Cx. quinquefasciatus* WGS data published in Haba et al. (2025), as described above. We then extracted the region which corresponded to our gene of interest plus 10,000 bp buffer on either side. Depth was then calculated using the -a flag within samtools and read depth and the fraction of sequenced specimens from each country containing the deletion were plotted in Rstudio. To test for evidence of the *GST-1* deletions, we allowed for two candidate deletion lengths which represented the deletion architecture we encountered in our sequencing data: a full-length deletion (breakpoints in chromosome NC_051862.1 at 160,428,836 and 160,429,887) and a shorter alternative deletion sharing the same start but ending at 160,429,282. For each individual, mean normalized depth was calculated in 10 bp windows immediately upstream and downstream of each of the three breakpoints (deletion start, full-length end and alternative end). An individual was classified as carrying the deletion if: (1) at the deletion start breakpoint, downstream depth was reduced by more than 15% relative to upstream depth (downstream mean < 0.85 × upstream mean), consistent with a drop in coverage entering the deletion; and (2) at either the full-length or alternative end breakpoint, downstream depth was elevated by more than 15% relative to upstream depth (downstream mean > 1.15 × upstream mean), consistent with a return to baseline coverage upon exiting the deletion. To confirm that the 10 bp windows were representative, we examined read depth in additional flanking windows and found it consistent with the genotypes assigned by the 10 bp approach. The per-population plots show the full-length deletion in red and the alternative deletion region in a darker shade to visualize the two boundaries simultaneously (Supplemental Figures 4-26). Our analysis considered only the presence or absence of the deletion and did not attempt to distinguish between heterozygous and homozygous genomes with the deletion.

### Manual genotype curation

For all specific SVs highlighted in the Results, genotypes called by Sniffles were inspected using the Integrative Genomics Viewer (IGV) [141,142]. Deletions were considered homozygous if zero reads spanned the breakpoints called in the VCF, and heterozygous if there were both reads spanning the deletion and reads supporting the breakpoints. Similarly, insertions were considered homozygous if there were no reads without the insertion that spanned the insertion breakpoint and considered heterozygous if some reads spanning the breakpoint contained the insertion and some did not. The predicted genotypes and manually curated genotypes for all SVs highlighted in the Results can be found in Supplemental Table 3.

## Results

### 1. High quality Culex quinquefasciatus genome assemblies from two population samples

We collected and sequenced the genomes of seven *Cx. quinquefasciatus* individuals from two locations, Puerto Rico and Zambia (Figure 1A). We generated an average of 60 Gb and 64 Gb of PacBio HiFi reads in Zambia and Puerto Rico samples, respectively. This resulted in an average sequencing depth that ranged from 67.7-83.6x in Zambia and 75.2-85.6x in Puerto Rico. With these sequences, we produced seven high-quality genome assemblies. Our genome size estimates ranged from 517-651 Mb with scaffold N50s ranging from 174 - 203 Mb (Table 1). Analyses using the Benchmarking Universal Single-Copy Orthologs program revealed that our assemblies exhibited a high degree of completeness that was similar to other mosquito genomes. We recovered 84.8-91.6% of the expected core genes with 7.1-13.5% of them missing across the seven individuals. To confirm our morphological species identification, we performed phylogenetic analyses using both mitochondrial and nuclear markers from our collections. Although our mitochondrial phylogeny lacked sufficient resolution to identify our samples down to the species level (i.e., our samples were indistinguishable from *Cx. quinquefasciatus*, *Cx. hutchinsoni* and *Cx. pipiens pipiens*; Supplemental Figure 27), our nuclear phylogenetic tree showed that both of our population samples formed a clade with the *Cx. quinquefasciatus* reference genome as opposed to any of the other seven *Culex* species we included in our analysis (Figure 1B).

**Figure 1.**
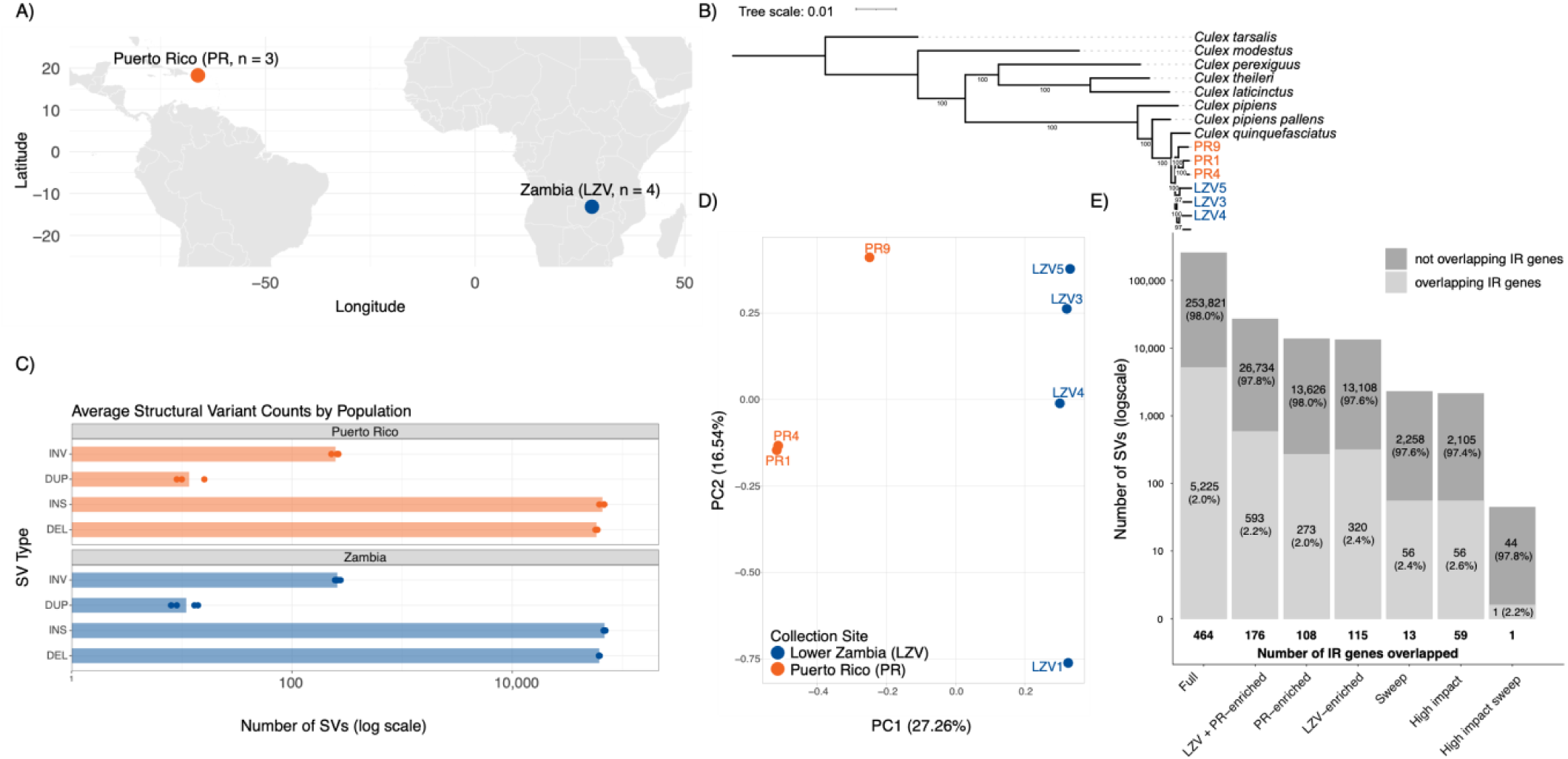
**A)** Map of sampling locations including the number of samples analyzed from each site. **B)** Phylogenetic relationships of our samples and eight published *Culex* reference genomes. Phylogeny was inferred from a multiple sequence alignment of the amino acid sequences of 2,893 orthologs that were present and single copy in 93.3% of samples under the JTT+F+R10 substitution model. Values at each branch represent bootstrap support and sample ID colors correspond to sampling locations. The tree was rooted using *Culex tarsalis* as the outgroup and visualized with iTOL. **C)** The average number of each type of structural variant split into the two sampling locations. The points indicate the counts for each individual. **D)** Principal Component Analysis plot based on structural variants for PC1 and PC2 with the percentage of total variance explained by each component. Samples are colored according to their sampling location. **E)** The number of structural variants that overlap insecticide resistance genes organized by the differently filtered sets of variants. This includes the full set, the population enriched sets (variants which have extreme allele frequencies between the two populations and the combination between these sets), the sweep set (variants which overlap selective sweeps), the high impact set (variants which are predicted to have high functional impacts by SnpEff), and the high impact sweep set (the intersection between the latter two sets). The number of insecticide resistance genes that are overlapped is in bold along the bottom of the plot (note that multiple SVs can overlap the same gene).

**Table 1.**
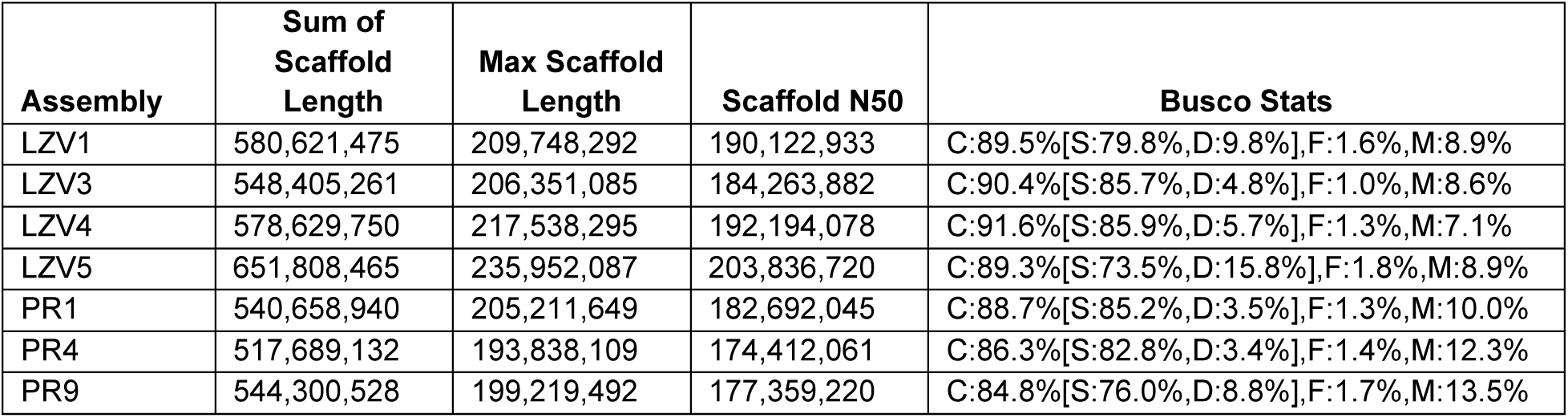
Final genome assembly summary statistics. BUSCO stats include complete (C), single-copy (S), fragmented (F), and missing (M).

| Assembly | Sum of Scaffold Length | Max Scaffold Length | Scaffold N50 | Busco Stats |
| --- | --- | --- | --- | --- |
| LZV1 | 580,621,475 | 209,748,292 | 190,122,933 | C:89.5%[S:79.8%,D:9.8%],F:1.6%,M:8.9% |
| LZV3 | 548,405,261 | 206,351,085 | 184,263,882 | C:90.4%[S:85.7%,D:4.8%],F:1.0%,M:8.6% |
| LZV4 | 578,629,750 | 217,538,295 | 192,194,078 | C:91.6%[S:85.9%,D:5.7%],F:1.3%,M:7.1% |
| LZV5 | 651,808,465 | 235,952,087 | 203,836,720 | C:89.3%[S:73.5%,D:15.8%],F:1.8%,M:8.9% |
| PR1 | 540,658,940 | 205,211,649 | 182,692,045 | C:88.7%[S:85.2%,D:3.5%],F:1.3%,M:10.0% |
| PR4 | 517,689,132 | 193,838,109 | 174,412,061 | C:86.3%[S:82.8%,D:3.4%],F:1.4%,M:12.3% |
| PR9 | 544,300,528 | 199,219,492 | 177,359,220 | C:84.8%[S:76.0%,D:8.8%],F:1.7%,M:13.5% |

### 2. Structural variants and single nucleotide polymorphisms reveal strong population differentiation between Zambia and Puerto Rico

After discovering and genotyping SVs using both read- and assembly-based methods and filtering for data quality as described in the Methods, our full set of SVs consisted of 259,046 variants: 115,002 deletions, 47 duplications, 929 inversions and 143,068 insertions (Supplemental Table 4). In all individuals, insertions were the most common SV type, followed by deletions, inversions, then duplications (Figure 1C, Supplemental Figure 28). We observed population stratification between the Zambian and Puerto Rican samples in each of our different analyses. In the phylogenetic tree, the two population samples form two distinct clades and both the SV-based PCA and SNP-based PCA show similar degrees of population structure. Both the SV-based (Figure 1D) and SNP-based PCA (Supplemental Figure 29) split the two populations along PC1. PC1 accounts for 27.26% and 36.7% of the variance in the data and PC2 accounts for 16.54% and 14.8%, respectively. Using SVs that were polymorphic within a given country, we generated the site frequency spectra for each population (Supplemental Figure 30) and show that they both have an excess of rare alleles.

### 3. Predicted loss-of-function effects are substantially more common among structural variants than single nucleotide polymorphisms

In addition to the 2,161 SVs that were annotated as ‘high impact’ by SnpEff (Supplemental Table 5 and 7), 473 SVs were annotated as ‘moderate,’ 522 as ‘low’ impact, and 258,549 as ‘modifiers’ (a classification which includes all changes outside the functional coding regions; Supplemental Figure 31). We further investigated the number of SVs that SnpEff classified as loss-of-function (LOF) alleles. For an SV to be considered LOF, we required that all transcripts of at least one gene were annotated with this effect. The full set of SVs contained 1,145 LOF SVs (0.44% of all SVs identified) across the seven samples. We also considered the number of LOF SVs in each individual genome—the seven genomes harbored between 502 and 568 LOF SVs each, with the number of genes affected ranging from 511 to 588 and the percentage of LOF SVs that were predicted as homozygous ranging from 32.0% to 39.2% in each sample (see Supplemental Table 6). 28 of the 1,145 LOF SVs from the full set were fixed across all seven samples, and 329 were present only in one sample, indicating that some of the apparent high-frequency LOF SVs could be low-frequency derived alleles, and vice versa. Furthermore, of the 1,039 total genes affected by LOF SVs, 604 were uncharacterized, which suggests that some of the affected genes could be non-coding or pseudogenes.

We also performed this analysis for the SNPs in our genomes and found that 2,552 of 40,017,054 SNPs (0.006%) in the multi-sample SNP dataset were predicted to induce LOF. The number of LOF SNPs in each genome ranged from 951 to 1,111, with 34.4 to 42.4% being homozygous, and these SNPs affected between 706 and 819 genes in each genome (Supplemental Table 6). 129 LOF SNPs were fixed across the seven samples, and 947 were present only in one sample. Of the 1,613 total genes affected by these SNPs, 918 were uncharacterized.

### 4. Structural variants are not enriched in selective sweep regions

To examine possible positive selection on SVs in our dataset, we ran SweepFinder2 using SNP data from three sub-Saharan African *Cx. quinquefasciatus* populations sequenced in Haba et al (2025) as these were the samples from Haba et al. that were located nearest to our Zambian population and thus may share some of the same SVs and selective pressures. We identified 1,331 CLR peaks across all populations and chromosomes (Figure 4 and Supplemental Figures 32-33). A total of 98 SVs from the LZV-enriched set overlapped a total of 80 peaks. We also found that two of the SVs from the LZV-enriched set that overlapped CLR peaks were annotated as high impact. One of these SVs was a 9,061 bp deletion that caused the complete loss of the lncRNA LOC119769733 and overlapped an intron of *neurobeachin-like protein 1* (LOC6035816). The other was a 5,168 bp deletion that caused the loss of LOC6032484, which has no annotated function but showed high amino acid sequence similarity to a histone-lysine N-methyltransferase 2C protein in *Cx. pipiens* via a BLAST analysis. We also identified 13 IR genes that contained or were within SVs that also overlapped selective sweeps (Figure 4; Supplementary Figures 32 and 33). A detailed analysis of all IR genes overlapping selective sweep windows can be found in the Supplementary Text. Following the procedure of our GO and IR gene enrichment analysis, we tested whether selective sweeps were enriched in SVs and found no significant enrichment ( *P-* value = 0.997).

### 5. Structural variants are enriched in insecticide resistance and sensory genes

We tested which GO terms were enriched within our structural variant datasets (full set, high impact, sweep set, high impact sweep, and the three population-enriched sets). Because each of the datasets was filtered to address a distinct biological question, GO enrichment analysis was performed independently for each dataset, and the results were interpreted within the context of the corresponding filtering strategy. After FDR correction, we found no statistically significant enrichment in any of our datasets except for the high impact set of SVs, where we found 28 GO terms that were significantly enriched (Supplemental Table 8). Of these 28 GO terms, we saw several terms involved in olfactory and chemosensory functions: odorant binding (GO:0005549), olfactory receptor activity (GO:0004984), sensory perception of smell (GO:0007608), and detection of chemical stimulus involved in sensory perception of smell (GO:0050911). There were also three GO terms which have been implicated in insecticide resistance, iron ion binding (GO:0005506), zinc ion binding (GO:0008270), and catalytic activity (GO:0003824) [87,143,144]. Lastly, additional enrichment terms were associated with pyruvate metabolism (GO:0004736), and proteolysis (GO:0006508).

To test whether genes previously implicated in insecticide resistance were overrepresented in the SV datasets, we used a list of 847 genes previously implicated in IR from the *An. gambiae* 1000 Genomes Project (Supplemental Table 1). Similar to our GO enrichment analysis, we ran this on each dataset independently. This gene list was generated prior to annotating genes within SVs so that the enrichment results would provide an unbiased assessment of whether IR genes disproportionately overlap SVs. We found that IR genes were significantly enriched in SVs in both the full set (*P*-value = 0.0019) and the high impact set of SVs (*P*-value = < 1/num_perms; Supplemental Table 9). In light of these findings, we investigated which IR genes and environmental sensing genes were affected by the SVs we identified.

Our seven SV datasets were examined for overlap with IR candidate genes (Figure 1E). In the full set of SVs, we identified 5,225 variants that intersected a total of 464 IR candidate genes. Within the high impact dataset, 56 SVs were annotated with high impact effects on 59 IR genes. The sweep dataset (which included variants that overlapped selective sweep regions identified by SweepFinder2) contained 56 SVs overlapping 13 IR genes, and the high impact sweep (which contained high SnpEff annotations and variants that overlapped selective sweep regions) dataset contained one SV overlapping an IR gene and its high impact SV (see complete list of overlapping genes in Supplemental Table 7). Our LZV-enriched dataset contained 320 SVs overlapping 115 IR genes, and the PR-enriched dataset contained 273 SVs overlapping 108 IR genes. The union of these sets contained 593 SVs that overlapped a total of 176 IR genes.

### 6A. Several structural variants overlapping insecticide resistance genes show evidence of population differentiation between Zambian and Puerto Rican populations

Within the LZV-enriched SVs, there were three insertions that intersected exons of IR genes. One of these variants impacted the coding sequence of *sorbitol dehydrogenase* (LOC6046345), via a 55 bp insertion within exon 1 of this putative IR gene (Supplemental Figure 34). This SV was predicted to cause a frameshift and loss of function. The ortholog of this gene in *Anopheles gambiae* identified by OrthoFinder is also annotated as *D-xylulose reductase A* [171] and is upregulated in deltamethrin-resistant strains of *An. funestus* [145]. Therefore, it may be surprising to find this variant segregating at a high frequency in our Zambian population given the prevalence of insecticide use in this region [88]. However, [146] showed that sorbitol dehydrogenase expression was downregulated in *Ae. aegypti* exposed to both high and low temperatures, implying a possible role for this gene in thermal adaptation. The insertion was genotyped by Sniffles as heterozygous in all Zambian samples except LZV5, which was homozygous, and absent from our Puerto Rican samples. Upon manual validation in IGV, we confirmed that these genotypes appear to be correct for all individuals aside from LZV5 and PR9, which both appear to be heterozygous. The fact that this SV is maintained at a frequency of 0.5 in Zambia indicates that it is unlikely to be highly deleterious, and its low frequency in the Puerto Rican population suggests that the deletion could be a source of local adaptation and/or balancing selection in Zambia; deeper sequencing would be required to investigate these possibilities.

There were two additional LZV-enriched SVs intersecting exons of IR genes—both insertions in the 3’ UTRs. One was a 128 bp insertion in the 3’ UTR of *cytochrome P450 9e2* (LOC6047797), which was genotyped as homozygous in LZV1 and LZV5 and heterozygous in LZV3 but appeared to be heterozygous in all three samples; the second was a 378 bp insertion in the 3’ UTR of *odorant receptor 82a* (LOC6043808), which was genotyped as homozygous in LZV1 and LZV3 and heterozygous in LZV5. Manual validation indicated that this SV was heterozygous in these samples, and also heterozygous in LZV4.

The Puerto Rico-enriched SVs contained five insertions and deletions overlapping exons, one of which impacted the coding sequence of an IR gene: a 385 bp insertion in exon 1 of the gene *glutaryl-CoA dehydrogenase, mitochondrial* (LOC6031211) on chromosome 2. Although this SV was genotyped as homozygous in PR1 and PR4 and heterozygous in PR9, manual inspection in IGV revealed that the variant was only present in PR9 and was indeed heterozygous in this sample. The remaining four SVs overlapped the 3’ and 5’ UTRs of IR genes. These included a 487 bp insertion in the 3’ UTR of *glutamate receptor ionotropic, kainate 2-like* (LOC119770850), which appeared to be heterozygous in LZV1, LZV5, and the Puerto Rican samples; a 74 bp insertion in the 5’ UTR of uncharacterized LOC6051012, which was heterozygous in LZV4 and absent in the other samples; and a 185 bp insertion in the 3’ UTR of *cuticular protein 49Ah* (LOC6042108), which was genotyped as homozygous in PR1 and PR4 but appeared heterozygous in both samples.

### 6B. High impact structural variants affect known insecticide resistance genes Glutathione S-transferases

Glutathione S-transferases (GSTs) are well-known insecticide resistance genes which impact metabolism and provide protection against oxidative stress following exposure to insecticides [38,147,148]. In our set of SVs predicted to have a high impact, we found a 1,051 bp exon deletion which encompasses the entirety of the first coding exon of the *GST1* gene (LOC6052515), probably resulting in complete gene inactivation in homozygotes (Figure 2A). This deletion was predicted to be heterozygous in LZV1 and LZV4, homozygous for the absence of the deletion in LZV3 and homozygous for the presence of the deletion in LZV5. All the samples from Puerto Rico were predicted to be homozygous for the reference. However further inspection within IGV revealed more complexity within this locus. First, LZV3 appeared heterozygous for the deletion. Second, PR1 and PR4’s reads do not support the presence of the 1,051 bp deletion, but instead suggest the presence of another complex SV which also appears to delete the first exon of the gene while inserting additional sequence nearby (Supplemental Figure 35). Analysis of this inserted sequence using BLAT [149] indicated that it had high sequence similarity to the transposable element DNA8-1_CQ. Due to *GST1* being a member of such a well-characterized IR gene family, we inspected the presence of this deletion in whole-genome short-read data sequenced by Haba et al. (2025). We plotted normalized read depth within this region and found that 21 of the 23 globally distributed population samples from Haba et al. had at least one individual appearing to contain the deletion, with several population samples containing multiple individuals with the deletion (Figure 2B).

**Figure 2.**
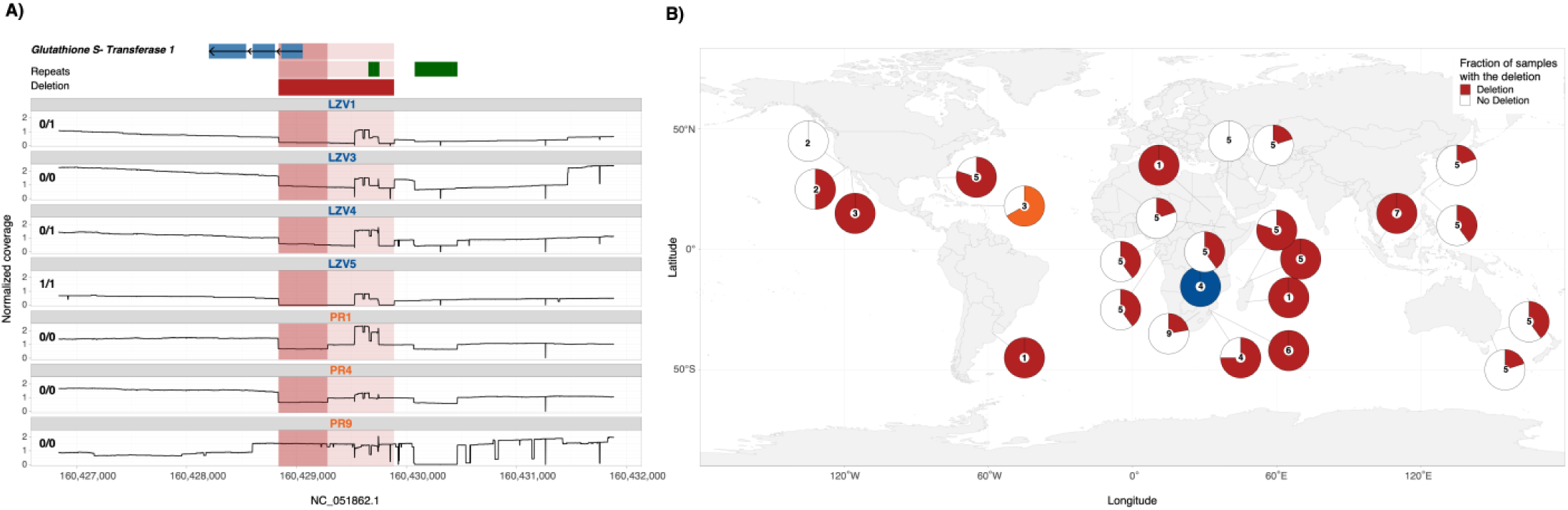
**A)** Normalized read depth plots for all seven individuals at a structural variant location that was predicted to have a high functional impact by SnpEff and overlapped an insecticide resistance gene, *Glutathione S-transferase 1*. Genotypes are recorded to the top left of the read depth plots, gene location and strand direction are in blue at the top, repeats are annotated in green, and the deletion is shown in red. The alternate deletion that is present in PR1 and PR4 is shown in darker red. **B)** Global prevalence of the deletion (including the alternate deletion) shown as pie charts with the number of samples shown at the center of the chart. These data were obtained from Haba et al. 2025 and the samples generated through this study are present in blue (Zambia) and orange (Puerto Rico).

#### Ionotropic receptors

Four of the high impact SVs overlapping IR genes affected ionotropic receptor genes. Insect ionotropic receptors are responsible for a variety of chemosensory functions, such as pheromone detection and temperature sensing [137–139]; but some ionotropic receptors, such as the GABA receptor encoded by the *Rdl* gene in *D. melanogaster*, are known insecticide targets [150,151]. By disrupting the function of these receptors, the SVs identified here could potentially serve as an environmental sensing or insecticide resistance modification. Two of the four high impact SVs affecting ionotropic receptors were specific to either the Zambian or Puerto Rican samples. This included a 118 bp deletion causing a frameshift in *ionotropic receptor 25a* (LOC6053318), which was heterozygous in PR1 and PR4 and absent in all other samples. Additionally, a 114 bp deletion in *ionotropic receptor 75a-like* (LOC119768765), which was heterozygous in LZV1 and LZV4, caused the loss of this gene’s stop codon.

We observed two additional high impact SVs in ionotropic receptor genes that were each only present in one sample, both of which were annotated as *ionotropic receptor 75a* (LOC6034998 and LOC119768764). Manual validation indicated that PR9 was heterozygous for an 83 bp insertion causing a frameshift and premature stop codon in LOC6034998, and LZV3 was heterozygous for a 184 bp deletion in LOC119768764. Finally, another gene which was annotated as *ionotropic receptor 75a,* but was not present in the IR gene list (LOC6044459), contained a 417 bp insertion in its fifth exon, which was predicted to cause a premature stop codon. This SV was heterozygous in LZV1 and LZV4.

#### Cytochrome P450s

Cytochrome P450s are the primary drivers of metabolic resistance to insecticides in many species [152]. These enzymes, which are involved in the catalytic oxidative metabolism of a broad range of compounds, typically confer resistance via gene duplication or overexpression [153–155]. Within our high impact set of SVs, there were 13 SVs impacting cytochrome P450 genes. Eight of these SVs occurred at low frequencies (present in only one of our 14 haploid samples, as confirmed through manual validation)—these included a 409 bp deletion removing the second exon of *probable cytochrome P450 304a1* (LOC6051151); a 272 bp deletion predicted to cause a gene fusion between two *probable cytochrome P450 9f2* genes (LOC6043062 and LOC6043064); a 230 bp insertion causing a frameshift and stop codon gain in exon 2 of *probable cytochrome P450 4ac1* (LOC6033127); a 71 bp insertion in exon 7 of LOC6049619 (annotated as *cytochrome P450 2A12/4A7*), causing a frameshift; a 197 bp deletion eliminating the start codon of *cytochrome P450 4C1* (LOC6049616); a 70 bp deletion overlapping exon 3 of LOC6042279 (annotated as *cytochrome P450 CYP4 family-like protein*), causing a frameshift; a 468 bp insertion causing gain of a stop codon in exon 9 of LOC6042285 (annotated as *cytochrome P450 4d10*); and a 2,270 bp deletion truncating the last exon of *probable cytochrome P450 28d1* (LOC6033808). Each of these low-frequency SVs, after manual validation, were determined to be heterozygous in a single sample.

Two of the high impact SVs overlapping cytochrome P450 genes were present at high frequencies in our samples. One of these was a 946 bp deletion in *probable cytochrome P450 6a14* (LOC6037494). The breakpoints of this deletion align with the boundaries of the gene’s first intron; additionally, these breakpoints align closely with a repeat region identified by WindowMasker [156]. Manual validation indicated that all Zambian samples were homozygous for this deletion, and all Puerto Rican samples were heterozygous. The second high-frequency SV was a 5,184 bp deletion that truncated exon 2 of another *probable cytochrome P450 6a14* (LOC6037496). This SV was manually verified as homozygous in all samples aside from LZV5, which was heterozygous. Due to the frequency of these SVs, we investigated them further by polarizing the alleles. We did this by mapping supporting reads to the outgroup species *Cx. pipiens* reference genome (GCA_963924435.1). When mapped to the *Cx. pipiens* reference, the SV-supporting reads did not exhibit a gap of the same size or at the same breakpoints as when mapped to *Cx. quinquefasciatus*. We therefore concluded that these SVs were insertions in the *Cx. quinquefasciatus* reference genome rather than deletions in our genomes (although we did observe evidence of an independent insertion in the *Cx. pipiens* reference genome).

We also identified three high impact SVs overlapping cytochrome P450 genes that occurred exclusively in the samples PR1 and PR4. These were a 64 bp deletion causing a frameshift in exon 3 of *probable cytochrome P450 28d1* (LOC6033808) (Supplemental Figure 36); a 433 bp deletion causing a frameshift in exon 3 of *probable cytochrome P450 9f2* (LOC6043064) (Figure 3A); and a 280 bp insertion causing a frameshift in exon 2 of probable cytochrome P450 6a13 (LOC6050716). All of these SVs were manually confirmed to be heterozygous in both PR1 and PR4.

**Figure 3.**
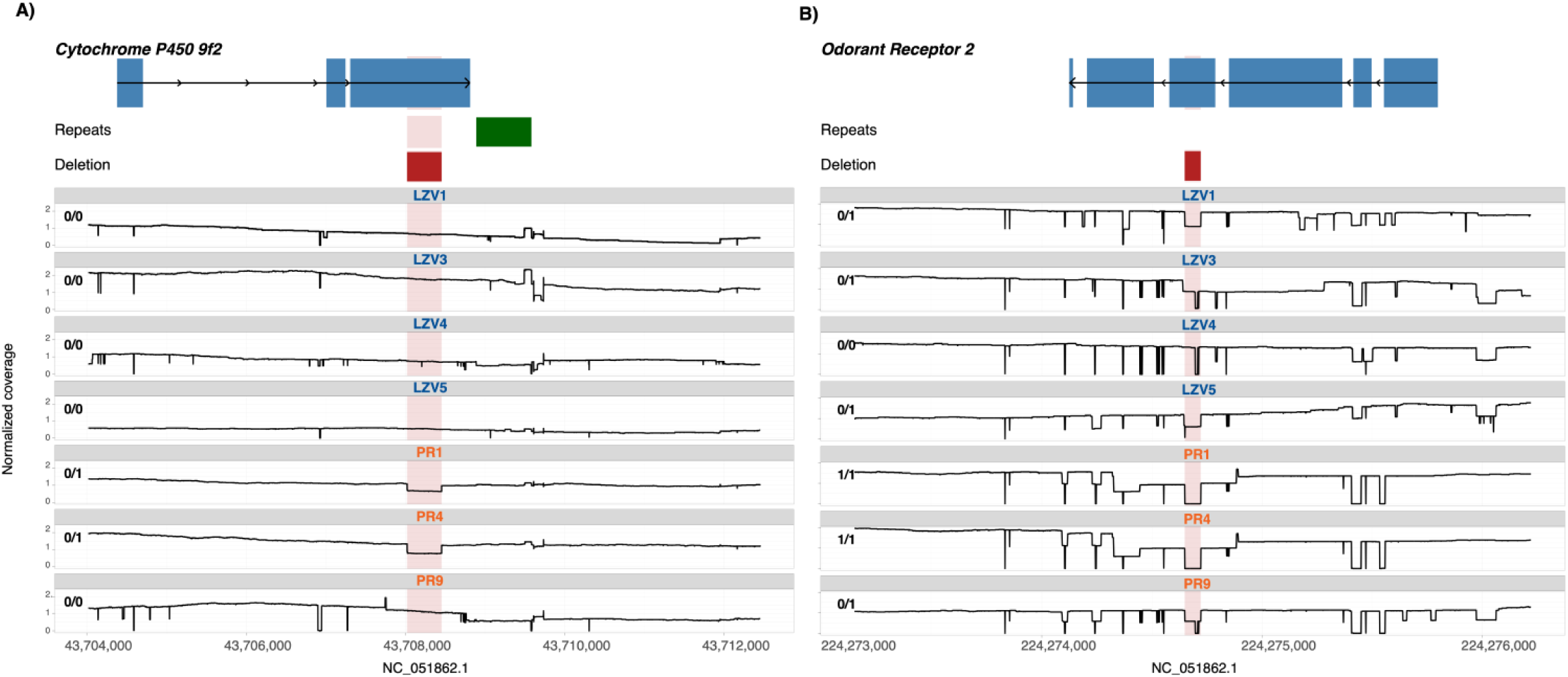
Normalized read depth plots for all seven individuals at structural variant locations that were predicted to have a high functional impact by SnpEff and overlapped an **A)** insecticide resistance gene, *cytochrome P450 9f2* and **B)** an environmental sensing gene, *odorant receptor 2.* Genotypes are recorded to the top left of the read depth plots, gene location and strand direction are in blue at the top, repeats are annotated in green, and the deletion is shown in red.

In addition to the cytochrome P450 genes in our IR candidate gene list, we also investigated the impact of SVs on *Cx. quinquefasciatus* genes annotated as cytochrome P450s that were not classified as orthologues of *An. gambiae* P450s implicated in IR. After repeating our procedure for identifying high impact SVs using these gene IDs, we found eight additional SVs that were predicted to have high impact effects on cytochrome P450 genes. Two were present in every sample: a 945 bp deletion in *probable cytochrome P450 6a14* (LOC6054219), which was manually verified as heterozygous in all samples except LZV5, which was homozygous, and a 242 bp deletion in LOC6054222 (annotated as *cytochrome P450 6B5*, *cytochrome P450 71B36*, and *cytochrome P450 CYP6 family-like protein*), which was homozygous in all samples. Both deletions had breakpoints aligning with intron boundaries, and supplementary alignments to the deleted regions indicated that the deletions possibly corresponded to repeats—therefore, these deletions could also represent the insertion of a repetitive element in the *Cx. quinquefasciatus* reference genome. The remaining high impact SVs affecting these cytochrome P450 genes occurred at lower frequencies.

#### Cuticle proteins

Cuticle thickening is a known mechanism of insecticide resistance that occurs due to overexpression of cuticle proteins, which prevents insecticide penetration [157,158]. The genes encoding these proteins are members of rapidly evolving gene families [159,160]. In our high impact set of SVs, there were three notable SVs that overlapped cuticle protein genes. One SV was a 436 bp insertion in the splice acceptor sequence of *larval cuticle protein A2B* (LOC6046702), which was heterozygous in PR1 and PR4 as confirmed in IGV. We also found a 4,105 bp deletion overlapping three cuticle genes: two *cuticle protein 8* genes (LOC6046729 and LOC6046730), and *cuticular protein 76Bd* (LOC6046728). This variant deleted the entire transcript of LOC6046729 and was predicted by SnpEff to cause a gene fusion between LOC6046730 and LOC6046728. This SV was heterozygous in LZV3, LZV4, and the Puerto Rican samples. We additionally observed a 728 bp deletion overlapping the 3’ UTR of *cuticular protein 64Ac* (LOC6035808), which was predicted as a gene fusion between *cuticular protein 64Ac* (LOC6035808) and *neurobeachin-like protein 1* (LOC6035816), as well as a splice region variant. This SV was manually confirmed as heterozygous in LZV3, LZV4, and LZV5, and was also found within a CLR peak (Figure 4). Finally, we identified an SV which had a predicted high impact on a cuticle protein gene that was not present in our IR gene list. This was a 9,762 bp deletion containing the entirety of *protein naked cuticle homolog* (LOC6042591), which was confirmed as homozygous in LZV1, LZV3, LZV4 and heterozygous in the remaining samples.

**Figure 4.**
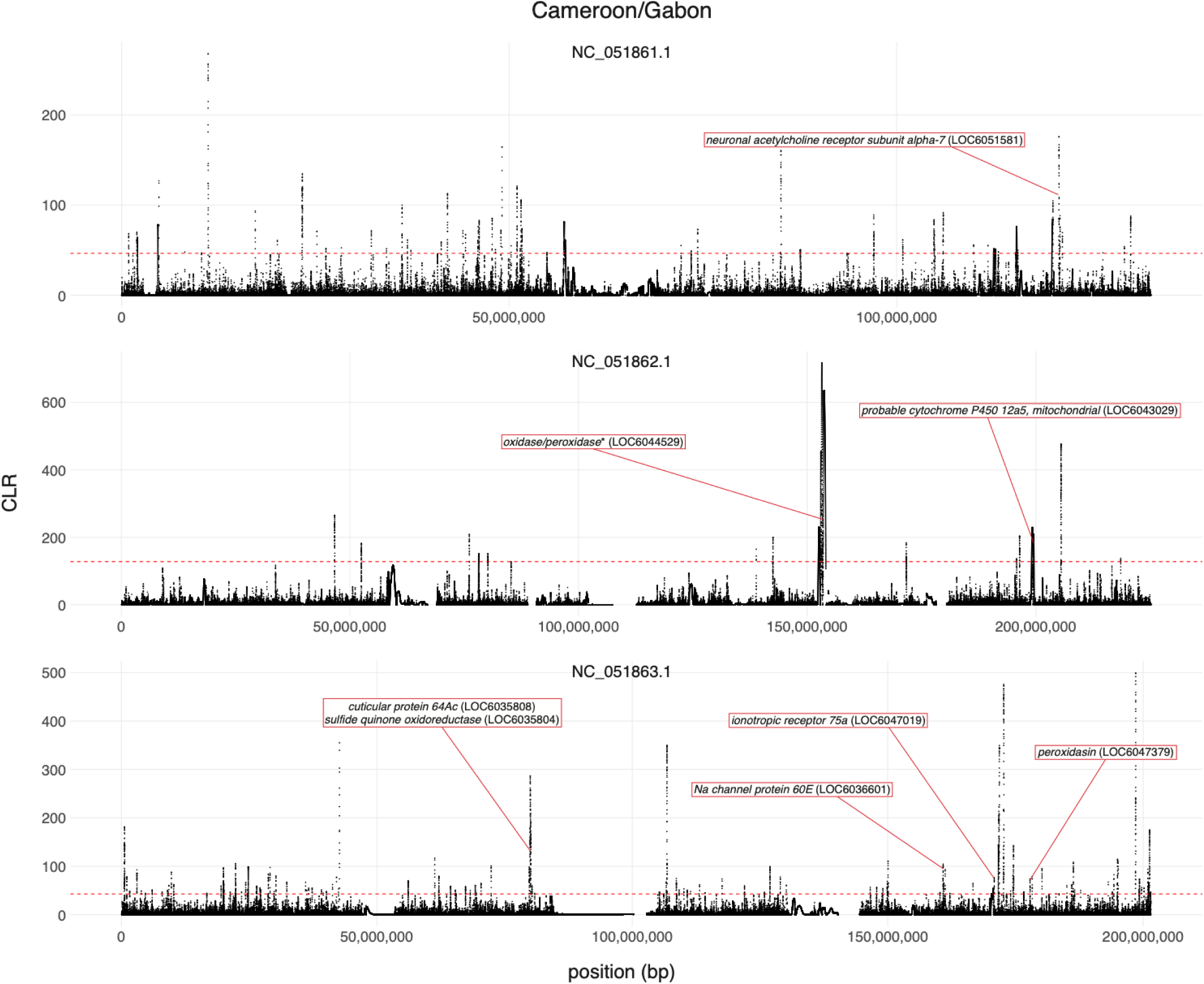
Candidate insecticide resistance genes overlapping both CLR peaks and structural variants in *Culex quinquefasciatus* populations from Cameroon and Gabon. Genes in red overlap SVs that are present in at least one individual from the Zambian population. SweepFinder analysis was performed using sequencing data from Haba et al (2025). The dashed red line indicates the 99^th^ percentile of CLR values across the chromosome, which was used to define a CLR peak. Genes with asterisks were annotated as uncharacterized loci by NCBI, but we included annotation information from the InterPro database here [174].

#### Untranslated regions

While SVs can exert effects through impacts on coding sequences, they have also been shown to alter gene expression through modifications to the untranslated regions of genes [161]. By examining our SnpEff-annotated VCF for SVs overlapping IR genes, we observed 95 SVs affecting the 3’ UTR of IR genes (annotated as “3_prime_UTR_variant”) and 32 SVs affecting the 5’ UTR of these genes (“5_prime_UTR_variant”). The 3’ UTR variants impacted 72 genes, and the 5’ UTR variants impacted 25 genes (Supplemental Table 8). It is possible that some of these genes’ expression levels are altered by their UTR SVs. Impacted genes spanned all four categories (metabolic, target site, behavioral, and cuticular; Supplemental Table 1). One of the 5’ UTR variants occurred at a high frequency and overlapped a SweepFinder peak—this was a 242 bp insertion in *sulfide quinone oxidoreductase* (LOC6035804), which was genotyped as homozygous in PR1 and PR4, and heterozygous in LZV4, LZV5, and PR9; however, manual validation suggests that this SV was not actually present in any Puerto Rican samples.

### 7. High impact structural variants affect environmental genes

In addition to assessing the impact of SVs on IR genes, we looked for overlap between SVs and genes with known roles in environmental response. Here, we highlight seven SVs with potential functional impacts selected from the high impact set of SVs. The first of these SVs was a 75 bp deletion overlapping exon 1 of *activator of 90 kDa heat shock protein ATPase homolog 1* (LOC119767536). This gene is responsible for activating a heat shock protein that maintains protein structure under heat stress [162], and the deletion was predicted to be a splice region variant. This SV was genotyped as heterozygous in LZV1, LZV3, and LZV5, homozygous in LZV4, and absent in the Puerto Rican samples; these genotypes were manually confirmed in IGV. We identified two high impact SVs in *pickpocket protein 28* (LOC6039638): a 191 bp deletion overlapping exon 2, which was annotated as a frameshift and splice donor variant, and a 371 bp deletion overlapping exons 4 and 5, which was annotated as a frameshift and stop codon loss variant. The 191 bp deletion was genotyped as heterozygous in LZV1, PR1, and PR4, homozygous in LZV5, and absent in the remaining samples. Manual validation confirmed each of these genotypes except for PR9, which was likely heterozygous for the SV. The 371 bp deletion was genotyped as heterozygous in LZV1, PR1, and PR4, and homozygous in LZV5; manual validation confirmed these genotypes. Pickpocket protein 28 is an ion channel responsible for gustatory water reception [163]. The gene *transient receptor potential cation channel protein painless* (LOC6037087) contained a 57 bp deletion overlapping exon 3, which was predicted as a frameshift and splice acceptor variant. The SV was genotyped as heterozygous in PR1 and PR4, which was confirmed in IGV. This gene is responsible for heat-induced pain response in *Drosophila* [164].

A 54 bp insertion in *protein lethal(2)essential for life* (*l(2)efl;* LOC6037263), which was present in all samples (manually validated as homozygous in LZV1, LZV3, LZV4, and PR1, and heterozygous in the others), was predicted to cause a premature stop codon in this gene. We also investigated this SV by polarizing the alleles against the *Cx. pipiens* reference genome—the reads containing the insertion mapped continuously to this genome, indicating that the insertion allele is likely the derived allele for this SV. In *A. aegypti*, *l(2)efl* was upregulated following dengue virus type 2 (DENV-2) infection and was shown to be a likely negative regulator of DENV-2 replication [165]. Further, in honeybees, *l(2)efl* expression was correlated with extreme desert temperatures common in the Arabian Peninsula [166]. We observed an 11,237 bp deletion affecting *heat shock protein 23-like* (LOC119770533), *protein lethal(2)essential for life* (LOC6037265), and *THAP domain-containing protein 2-like* (LOC119770528). Both LOC119770533 and LOC6037265 encode heat shock proteins which contribute to heat and cold tolerance [167,168]. This SV, which was manually confirmed as heterozygous in LZV3, PR1, and PR4, resulted in the complete deletion of all three genes. Finally, a 591 bp insertion in the first exon of *phenoloxidase-activating factor 1* (LOC6035180), which was predicted to cause a stop codon gain, was genotyped as homozygous in LZV5, heterozygous in the remaining Zambian samples, and absent in the Puerto Rican samples (Supplemental Figure 37). However, manual validation indicated that the insertion appeared to be heterozygous in LZV5, PR4, and PR9, and absent in the remaining samples. Phenoloxidase contributes to insect immunity by activating the melanization cascade in response to pathogens [169,170].

We identified nine high impact SVs that impacted odorant receptor genes, several of which segregated at high frequencies within our samples. Odorant receptors are a rapidly evolving gene family in insect species [136,171,172]. Their primary role is in environmental sensing, such as host seeking [136]; however, there is also evidence to suggest that these receptors can contribute to insecticide resistance by mediating behavioral avoidance of insecticides [173]. Three of these SVs were present in both Puerto Rico and Zambia. These include a 230 bp insertion in exon 1 of *odorant receptor 67a* (LOC6035727), which was predicted to cause a frameshift. All seven of our samples were homozygous for this insertion, which was manually validated as 172 bp in length. However, by polarizing this allele against the *Cx. pipiens* genome, we confirmed that this SV is likely a deletion in the *Cx. quinquefasciatus* reference genome. Another frameshift-causing SV, a 73 bp deletion in exon 4 of *odorant receptor Or2* (LOC6047488), was present in all samples excluding LZV3 and LZV4 (Figure 3B)—it was homozygous in PR1 and PR4, and heterozygous in the remaining samples (LZV3 was genotyped as heterozygous in the VCF, but was manually confirmed as heterozygous for a separate, larger deletion allele). Finally, we identified a 513 bp deletion in an intron of *odorant receptor co-receptor* (LOC6042040), which was predicted to cause a feature ablation and gene fusion with *transfer RNA alanine (anticodon AGC)* (LOC119771000). After manual validation, we determined that this deletion was homozygous in PR9 and heterozygous in all remaining individuals.

Three of the high impact SVs overlapping odorant receptor genes were unique to the Puerto Rican population. All three were heterozygous in PR1 and PR4 and absent in all other samples. These included: a 185 bp insertion in exon 1 of LOC6035750, which has a predicted odorant binding function [174,175], causing a frameshift and premature stop codon; a 172 bp deletion overlapping exon 2 of LOC119766993, which also had a predicted odorant binding function [174,175], which caused loss of the start codon; and a 654 bp deletion overlapping the 3’ UTR of LOC6035760, annotated as odorant receptor 36 [176], which SnpEff annotated as a splice region variant and bidirectional gene fusion with LOC6052980. However, the deletion is contained entirely within intronic sequence of LOC6052980, so it is unlikely to have a functional impact on this gene.

Finally, three of these nine SVs were present in only one of our samples. These were a 209 bp insertion in exon 2 of LOC6040695 (annotated as odorant receptor 9a) [177] which was heterozygous in LZV4, a 71 bp deletion in exon 2 of *odorant receptor 94a* (LOC6045705), which was heterozygous in LZV1, and a 737 bp insertion in exon 3 of *odorant receptor Or2* (LOC6047488), which was heterozygous in LZV1. Each of these SVs was predicted to affect their respective gene via a frameshift, with the 737 bp insertion also introducing a premature stop codon. We also identified additional SVs affecting odorant receptor genes which were present in our high impact sweep set of SVs—these included a 199 bp deletion causing the loss of a start codon in *odorant receptor 7a* (LOC6052036), which was heterozygous in LZV1, LZV5, PR4, and PR9; and a 233 bp deletion in exon 1 of *putative odorant receptor 83c* (LOC6036946) causing a frameshift, which was genotyped as heterozygous in LZV1, LZV3, PR1, and PR4, but appeared heterozygous in LZV3 only according to our manual validation.

## Discussion

Understanding the genetic basis of adaptation to anthropogenic stressors is a crucial and timely goal in evolutionary biology, with structural variants (SVs) likely playing a key role [178–180]. In this study, we uncover the widespread nature of SVs in the genomes of a major disease vector, *Culex quinquefasciatus*. We show that the composition of SVs largely reflects population structure and highlight SVs in genes associated with rapidly evolving phenotypes, such as those related to insecticide resistance and environmental sensing. We reveal many shared and population-specific SVs, SVs that have high functional downstream impacts, and SVs that may be targets of positive selection. Further, we discover an SV that appears to inactivate a gene in a well-known IR gene family (glutathione S-transferases) and that has a near global distribution. Taken together, these findings suggest that SVs may be implicated in rapidly evolving phenotypes in vector species more than previously appreciated and should therefore be considered alongside other forms of genetic variation known to contribute to IR. Although classical target-site resistance is most often associated with nonsynonymous mutations (e.g., *kdr* [181–183] or *rdl* [151,184,185]), emerging evidence suggests that structural variation at these loci may also contribute to IR [42,85,186–190]. More broadly, SVs may contribute to IR through changes in gene dosage, gene disruption, or alterations to regulatory architecture, however these mechanisms remain largely overlooked by conventional SNP-based approaches. Consequently, the relative paucity of demonstrated associations between SVs and IR should therefore be interpreted cautiously, as it may reflect the limited application of SV-sensitive approaches applied to insect vector and pest populations.

The use of insecticides can drive rapid evolutionary change by eliminating a substantial proportion of the population within a short period of time, allowing variants that confer resistance to quickly rise in frequency [40,191,192]. We sampled populations from Zambia and Puerto Rico, both of which have used insecticides to control mosquito populations, although insecticide use has likely been more extensive in Zambia [193,194]. Zambia’s malaria control campaign has used pyrethroids and organophosphates, as well as both long-lasting insecticide-treated nets (LLINs) and annual indoor residential spraying (IRS) [195]. In Puerto Rico, aerial spraying of organophosphates and ultra-low volume applications of insecticides have been used in government campaigns to combat Zika virus and dengue [86,196]. These control efforts would be expected to impose a strong selective pressure on the *Cx. quinquefasciatus* populations studied here, motivating investigation into the genomic features that might fuel the evolution of IR. In the present study, we found hundreds of IR genes overlapping SVs which were predominantly composed of insertions and deletions. They encompassed many different classes of possible IR genes, including glutathione S-transferases, cytochrome P450s, ionotropic receptors, and cuticle genes.

In *Culex* species, and mosquitoes as a whole, there has been a focus on investigating duplications in known IR genes [84,85,197–199] as copy number variants have been shown to contribute to IR through amplified expression [200] or neo-functionalization of new gene duplicates [201]. Indeed, gene duplications in cytochrome P450s [81], the voltage-gated sodium channel [42], and the *ace-1* gene [85,202,203] have been reported in insecticide resistant strains of *Culex* species, with the latter also showing evidence of common duplications in *D. melanogaster* [204–206]. However, there has been far less attention on other SV types, despite their potential adaptive contribution [178]. Here we found that small-to medium-sized deletions and insertions were much more prevalent than duplications and inversions and often appeared likely to have high functional downstream impacts. Indeed, 1,043 of our 2,161 high impact SVs were smaller than 1,000 bp and ∼99% of these SVs were insertions and deletions. When considering SVs larger than 100,000 bp, the total number of high impact SVs only increased by 79 additional variants, with insertions and deletions smaller than 1,000 bp still representing >95% of the high impact variants. Given the disproportionately high downstream functional impacts of these types and sizes of SVs, future studies should seek to incorporate them whenever feasible.

Loss-of-function mutations or even complete losses of IR target genes are a well-documented route for adapting to insecticide exposure. In *Drosophila,* the *Met* mutation, which eliminates the insecticide target protein altogether, has been shown to increase rates of IR (but see; [207])[208,209]. However, this is not an expected form of resistance in metabolic resistance genes where elevated expression levels tend to be associated with increased IR [210,211]. Many of the SVs we describe here overlap or completely contain IR metabolic resistance genes. For example, we detected two deletions that overlap exons of cytochrome P450 genes *9f2* and *28d1* that were predicted to be loss of function mutations and were present in two of our seven samples (PR1 and PR4). Similarly, we detected two separate deletions in the same *glutathione S-transferase 1* gene with the larger of the two deletions occurring in each of the Zambian samples and the smaller deletion only occurring in two of the three Puerto Rico samples. Both of these deletions encompass the entire first coding exon, likely indicating a loss-of-function mutation in homozygotes. Given that glutathione S-transferases are key metabolic genes involved in xenobiotic detoxification [212], one would expect a loss-of-function mutation to be deleterious. However, when we explored the prevalence of these deletions in global whole genome sequencing [124], we found that one or the other was present in 21 of the 23 populations sampled and that 51% of all individuals sampled have the deletion in either heterozygous or homozygous form, indicating that in at least some environments it is not strongly deleterious. Future work will be required to determine whether deletions of this gene have been positively selected in some subpopulations of *Cx. quinquefasciatus*.

Evolutionary trade-off theory may help to explain why deletions in detoxification enzymes may not be deleterious. One possibility is that resistance mechanisms impose energetic or physiological costs, such that reduced insecticide exposure relaxes selection for maintaining costly resistance traits [213]. Indeed, in *Cx. pipiens*, increased insecticide resistance was shown to reduce energetic reserves by 30% on average [214] and in *D. melanogaster*, increased resistance levels were associated with lower overwintering success [215]. Under these circumstances, deletions or downregulation of resistance-associated genes may be favored when the selective pressure imposed by insecticides is reduced. Given that many of the SVs highlighted here are deletions specific to the Puerto Rican population, this excess of SVs in IR genes may reflect relaxed negative selection in this population. Weaker negative selection would allow for an increase in the accumulation of SVs relative to genes subject to stronger selective constraints. While it is tempting to speculate that these deletions in Puerto Rico may be attributable to potentially lower rates of insecticide use compared with Zambia (although direct comparisons are difficult), our enrichment analysis did not find clear support for this interpretation, as we did not see population-specific evidence of enrichment for IR-related functions or genes. It is important to note that even in the absence of insecticides, we may not expect IR genes to be under weaker selection than other genes given that many may perform biological roles outside of IR (e.g. [216]). An alternative explanation that we cannot rule out is that the usage of different insecticides in different locations could potentially result in a trade-off where resistance to one insecticide confers sensitivity to the other, a phenomenon known as negative cross-resistance. This has been demonstrated in *Anopheles* where over-expression of *Cyp6* increased susceptibility to malathion while reducing susceptibility to pyrethroids [217]. Further studies are needed to distinguish between these scenarios and investigate the phenotypic impact of the SVs.

Although SVs are typically expected to be more deleterious than SNPs [218], they also may potentially be a source of large-effect adaptations in populations challenged by severe selective pressures, such as insecticide exposure [49,219]. Interestingly, we found significant enrichment of IR genes in two of our datasets and several significantly enriched GO terms that have been previously implicated in IR. These included iron and zinc binding and catalytic activity [87,143,144]. Importantly, enrichment of IR genes and related GO terms alone would not necessarily provide evidence of adaptation, as such patterns could instead indicate mutational bias towards IR genes, selection on traits that are correlated with IR, or stabilizing selection. To further investigate whether SVs in our dataset may be subject to positive selection, we ran SweepFinder on three representative populations from Haba et al. (2025) that face similar selective pressures to our Zambian population. In our Zambia-enriched SVs, we found 98 SVs that overlapped CLR peaks identified by SweepFinder. Two of these SVs were annotated as high impact by SnpEff, and although both impacted genes were uncharacterized, the presence of a sweep overlapping the SVs suggests that these variants could induce a phenotype that is the target of positive selection (e.g., insecticide resistance). Together, these findings indicate that, while we found no evidence for genome-wide enrichment of known IR genes in this set of SVs, a subset of Zambia-enriched SVs may nonetheless have adaptive significance and warrant further investigation.

Olfactory receptor genes and immune genes evolve rapidly because they interact directly with an everchanging external environment full of pathogens and chemical cues [136,220]. Both categories of genes are statistically enriched for gene duplications in *Hermetia illucens* (the black soldier fly), relative to the most recent common ancestor node of Stratiomyidae [221]. Insect odorant receptors (ORs) form a large and divergent gene family that represents a radiation exclusive to insects [222]. This gene family enables insects to smell and taste chemicals in their environment and the large radiation combined with dramatic differences in gene family size between species has led to the belief that OR genes may evolve rapidly [136,220]. Here, we found many SVs which overlapped genes that are involved in sensory perception of smell. Specifically, we saw nine high impact SVs which contained or overlapped odorant receptor genes and two which were high impact and overlapped CLR peaks which may indicate that they are under selection. One of these genes, *Or2*, was shown to be highly sensitive to the oviposition attractant indole in *Cx. quinquefasciatus* [223]. Further, our high impact SVs were significantly enriched for olfactory and chemosensing functions, suggesting that SVs may be important contributors to variation in these phenotypes.

We also describe an insertion in the innate immune gene *phenoloxidase activating factor 1* that is a component of the pro-phenoloxidase activating system [224]. Invertebrate phenoloxidase is one of the most important tools against pathogens and is involved in the melanization of pathogens [224,225]. Phenoloxidase was also shown to have genetic correlations with costly functions where, for example, selection on phenoloxidase activity in yellow dung flies led to earlier adult mortality when there was no available food [226]. The loss-of-function insertion is found in Zambian individuals and is frequently observed in the heterozygous state. Given the important role of this immune gene, its population-specific distribution may reflect differences in environmental pathogen exposure, whereby loss of gene function may be advantageous under pathogen pressures specific to the Zambian environment. However, we do not see significant enrichment for immune-related functions in any of our SV datasets, suggesting that this specific SV may not be representative of a broader enrichment of structural variation affecting immune function.

Although it is clear that SVs are involved in a wide range of biological functions ranging from IR to environmental sensing and response, there are several key caveats that need to be acknowledged. First, although our results suggest that the functional effects of many of our SVs are considerable, this prediction is based on genomic annotations and thus the exact phenotypic impact of these SVs is unknown. Second, while it is tempting to assume that an SV that overlaps a selective sweep is in fact the target of the sweep, it may have simply hitchhiked with the sweeping allele. For example, we observed sweep windows overlapping SVs present at low to intermediate frequencies in our Zambian population, such as the high impact deletion in *odorant receptor 7a* which was heterozygous in two Zambian samples. The presence of a selective sweep intersecting this low-frequency SV is consistent with a deleterious variant reaching higher frequency than it otherwise would be expected to because it is linked to a positively selected allele [227,228]; we also again emphasize that because we have not systematically polarized our SVs, apparent low-frequency alleles could be high-frequency derived alleles and vice versa. Finally, although some of the SVs described here may be adaptive, others may be deleterious or may be primarily evolving under neutral processes.

This study is the first to use long-read sequencing to study populations of *Cx. quinquefasciatus*, and in doing so, we reveal a large number of SVs (>259,000) within this species’ genome. The number of SVs reported here far exceeds the number seen in *Drosophila* species with estimates ranging from an average of approximately 3,000 across 5 lines (each of which contained 100-200 individuals; [229]) to 20,000 across 14 genomes [49]. While different methodology, a highly diverged reference genome, or false positives and negatives may contribute to the difference in the number of SVs observed here, it is unlikely to account for the >13-fold increase in the number of reported SVs. To test the potential contribution of the latter two scenarios, we excluded SVs where the non-reference allele was fixed in all of our samples and SVs where the minor allele was present in only one individual and we were left with a total of 255,546 and 216,541 SVs, respectively. Although false positives may not always appear as singletons, we conducted extensive manual curation of specific SVs and found that, while the genotypes did not always appear accurate, evidence for the existence of the SV typically appeared to be sound. Therefore, it does not appear that large numbers of variants specific to the reference genome or putative false positives can explain this difference alone. Unfortunately, no studies have used long-read sequencing to investigate SVs in mosquito species and so we lack a sufficient comparison. A recent study using short-read sequencing in *An. stephensi* reported approximately 3,000 duplications and 16,000 deletions across 115 genomes [230], markedly fewer than the numbers reported in this study. We therefore posit that the stark difference in the number of SVs in *Cx. quinquefasciatus* compared to that of both *Drosophila* and *Anopheles* is instead a result of the combination of different methodologies and genomes which contain the genetic substrate required for the proliferation of genomic rearrangements, namely the abundance of TEs and repetitive sequences. These findings highlight the critical importance of expanding the application of long-read sequencing to include other vector species. Finally, given the extensive SV landscape, the outsized functional impact of SVs vs SNPs (0.44% of all SVs were predicted to result in loss of function compared to 0.006% for SNPs) and the fact that ∼5,000 of our SVs overlap nearly 500 IR genes, it is clear that long-read sequencing is a vital yet underutilized approach for uncovering genetic variation affecting key phenotypes in these important disease vectors.

## Acknowledgments

We would like to thank Dr. Edward G. Smith for his helpful discussions and edits to the manuscript.

## Data availability

All sequencing data used in this paper will be made publicly available prior to publication. All code used for the analyses presented in this paper can be found at https://github.com/SchriderLab/SVs_Culex_quinquefasciatus_2026.

