## Supplemental Documents for "Long-read sequencing reveals genomic landscape of structural variants in a major disease vector"

### Supplemental Figures

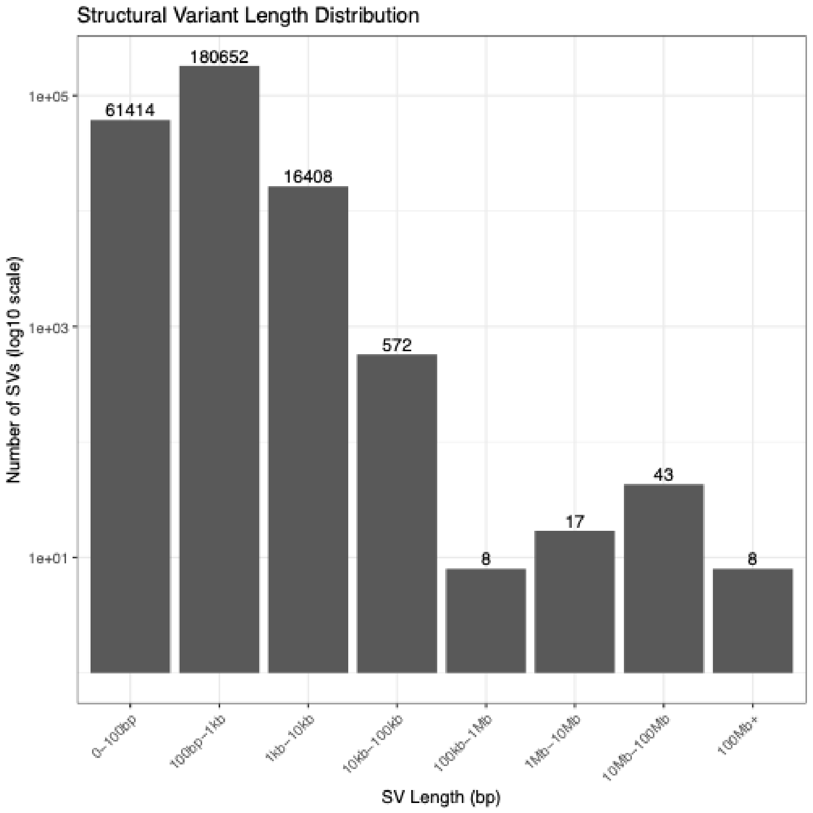


**Supplemental Figure 1.** Distribution of structural variant lengths in the genotyped and merged VCF. Based on the increase in SV count above 1 Mb and manual inspection of read depth within and surrounding the SVs 100 kb-1 Mb in length, which indicated that the SV calls within this length range were false positives, we filtered out SVs >100 kb for our final dataset (the "full set" of SVs).

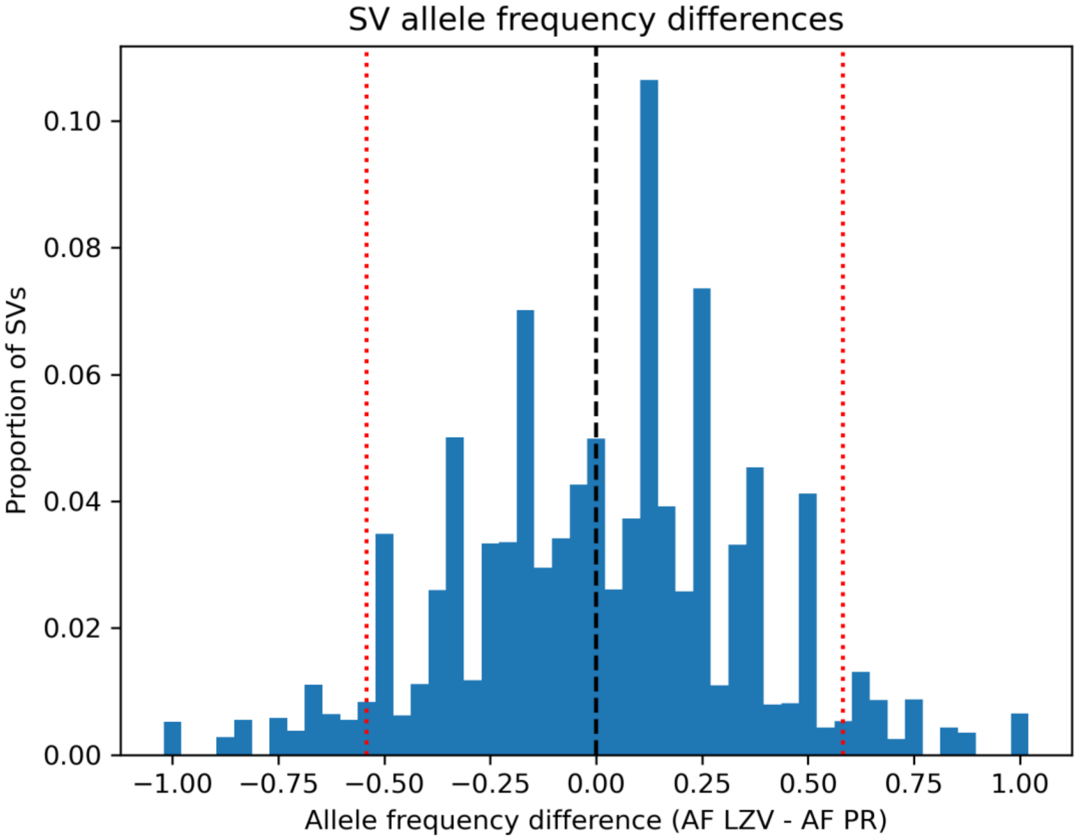


**Supplemental Figure 2.** Distribution of SV allele frequency differences between the Zambian and Puerto Rican populations. This distribution was used to define cutoffs for our LZV-enriched and PR-enriched set of SVs: the LZV-enriched VCF contained SVs in the top 5% of allele frequency differences between LZV and PR, and the PR-enriched VCF contained SVs in the bottom 5%. The top and bottom 5% is shown with the dotted red line.

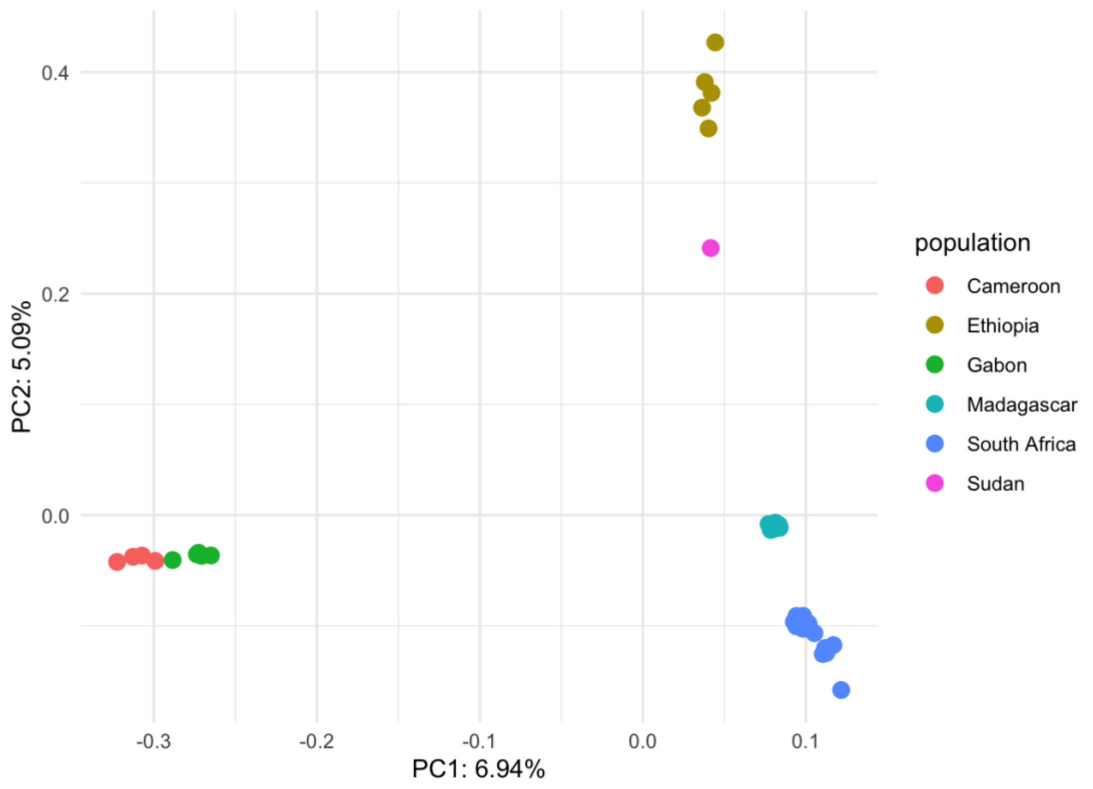


**Supplemental Figure 3.** PCA of SNPs in sub-Saharan African *Cx. quinquefasciatus* populations from Haba et al (2025). Our goal was to identify population samples that may be experiencing similar selective pressures to our Zambian population and investigate signatures of selection in these samples. Based on this clustering and the sample sizes of each population, we used three populations for our SweepFinder analysis: South Africa, Madagascar, and a combined population comprised of the individuals from Cameroon and Gabon.

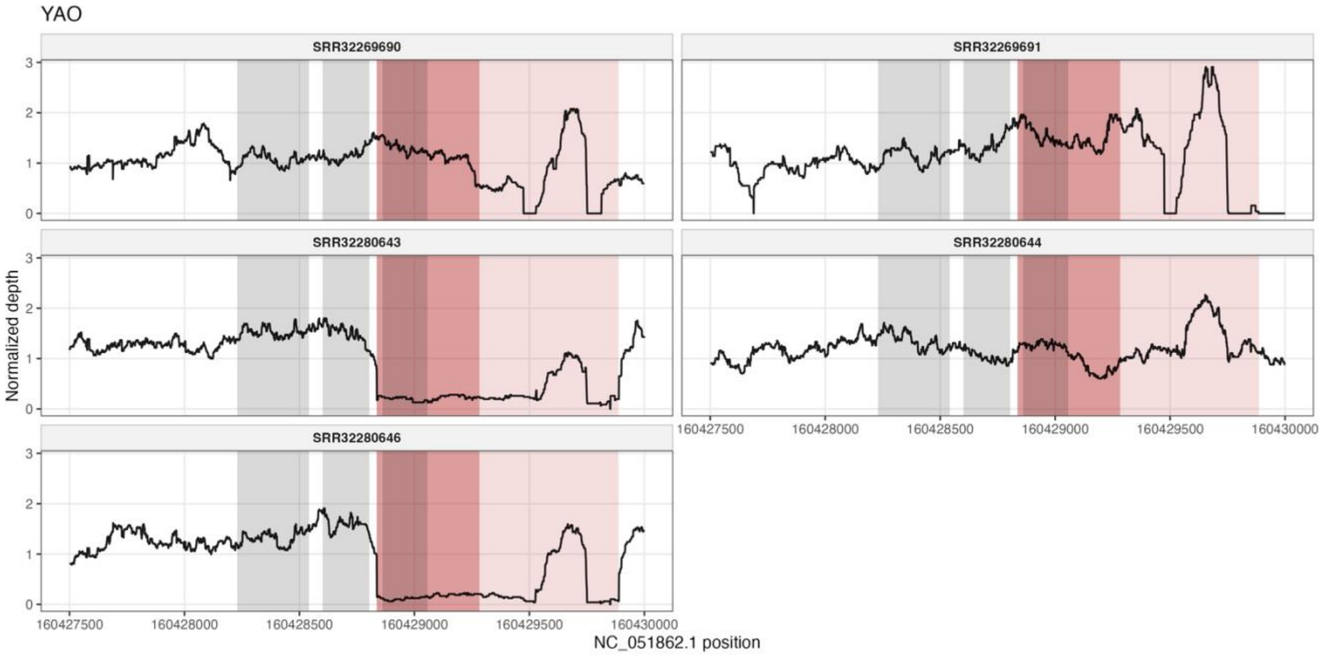


**Supplemental Figure 4:** Normalized read depth plots for the *GST-1* deletion region in each of five individuals from Yaounde, Cameroon. Read depth was normalized by that individuals average read depth across the entire genome. Data was obtained from Haba et al. 2025. Light red shaded regions indicate the larger deletion while darker shaded region indicates the deletion with an alternative ending and grey shaded regions indicate each exon in the *GST-1* gene.

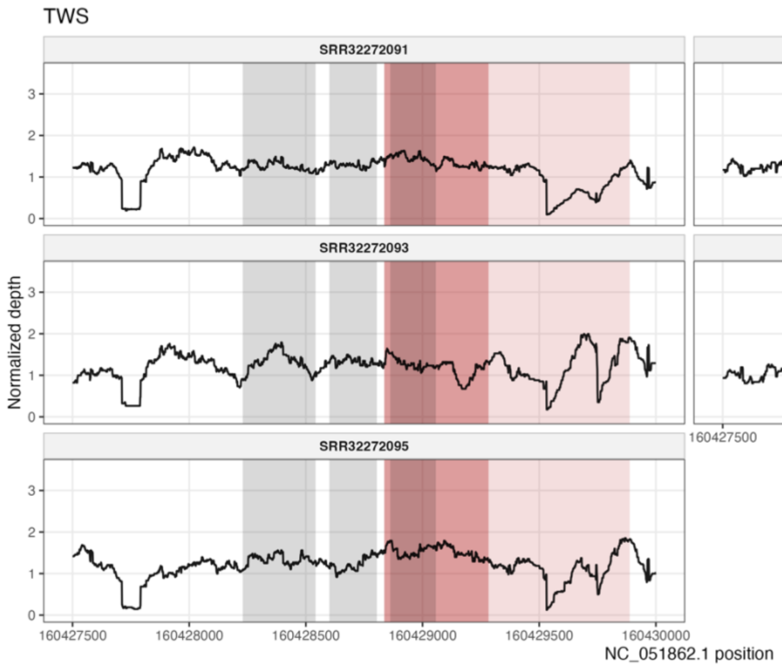


**Supplemental Figure 5:** Normalized read depth plots for the *GST-1* deletion region in each of five individuals from Chiayi County, Taiwan. Read depth was normalized by that individuals average read depth across the entire genome. Data was obtained from Haba et al. 2025. Light red shaded regions indicate the larger deletion while darker shaded region indicates the deletion with an alternative ending and grey shaded regions indicate each exon in the *GST-1* gene.

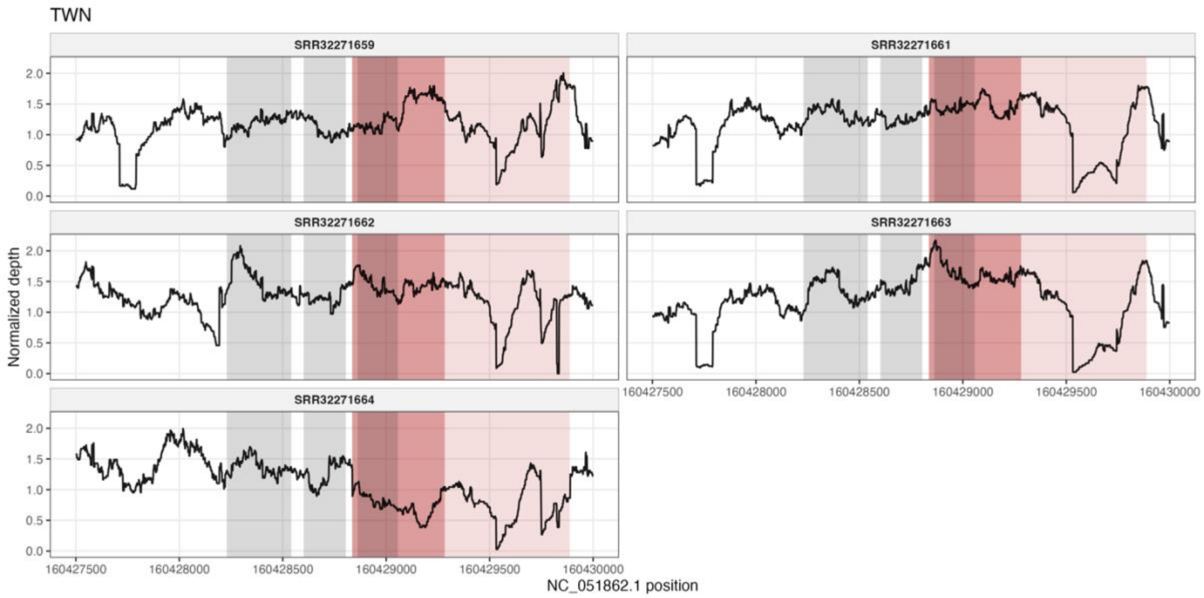


**Supplemental Figure 6:** Normalized read depth plots for the *GST-1* deletion region in each of five individuals from Taipei, Taiwan. Read depth was normalized by that individuals average read depth across the entire genome. Data was obtained from Haba et al. 2025. Light red shaded regions indicate the larger deletion while darker shaded region indicates the deletion with an alternative ending and grey shaded regions indicate each exon in the *GST-1* gene.

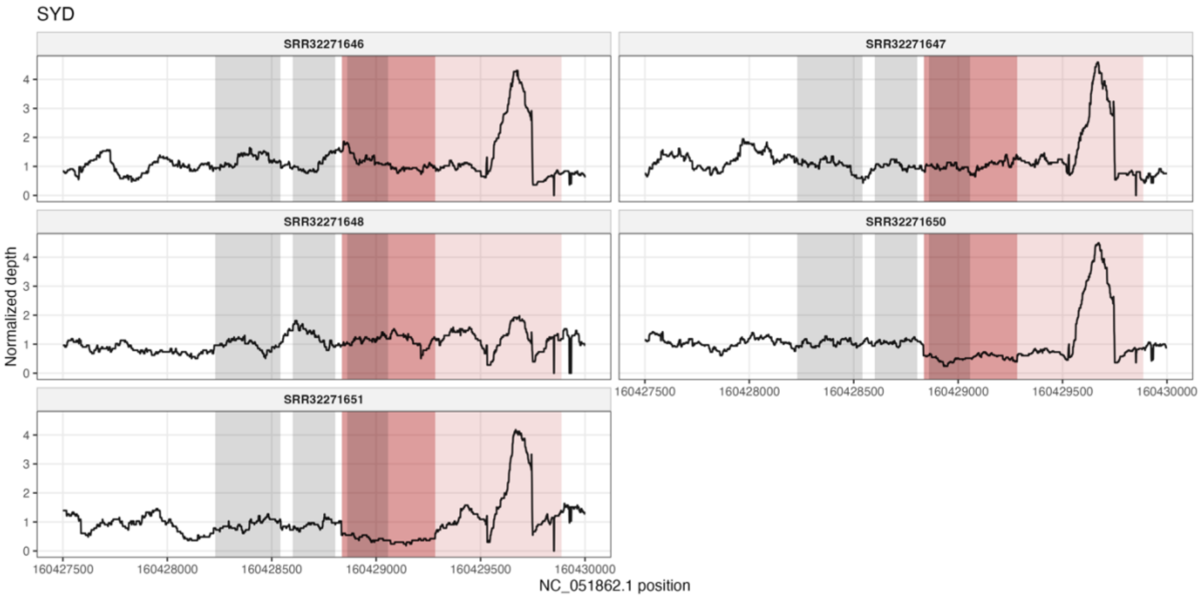


**Supplemental Figure 7:** Normalized read depth plots for the *GST-1* deletion region in each of five individuals from Newington, Australia. Read depth was normalized by that individuals average read depth across the entire genome. Data was obtained from Haba et al. 2025. Light red shaded regions indicate the larger deletion while darker shaded region indicates the deletion with an alternative ending and grey shaded regions indicate each exon in the *GST-1* gene.

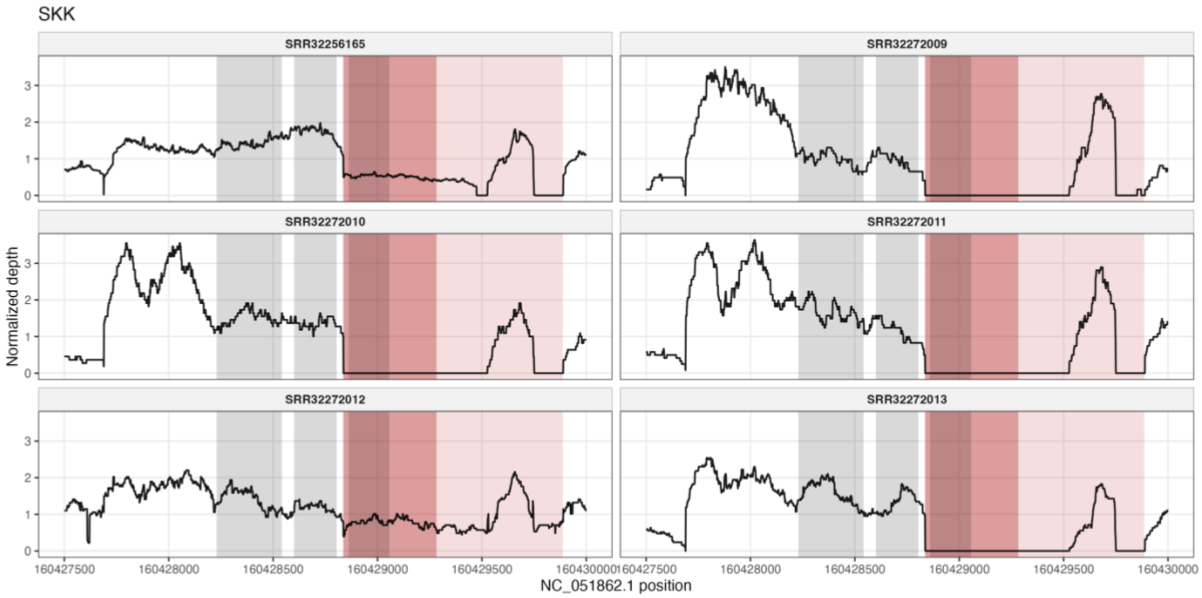


**Supplemental Figure 8:** Normalized read depth plots for the *GST-1* deletion region in each of six individuals from Skukuza, South Africa. Read depth was normalized by that individuals average read depth across the entire genome. Data was obtained from Haba et al. 2025. Light red shaded regions indicate the larger deletion while darker shaded region indicates the deletion with an alternative ending and grey shaded regions indicate each exon in the *GST-1* gene. Note that we collapsed samples collected from Skukuza (SRR32272009-13) with those collected from Mpumalanga (SRR32256165) due to their geographic proximity.

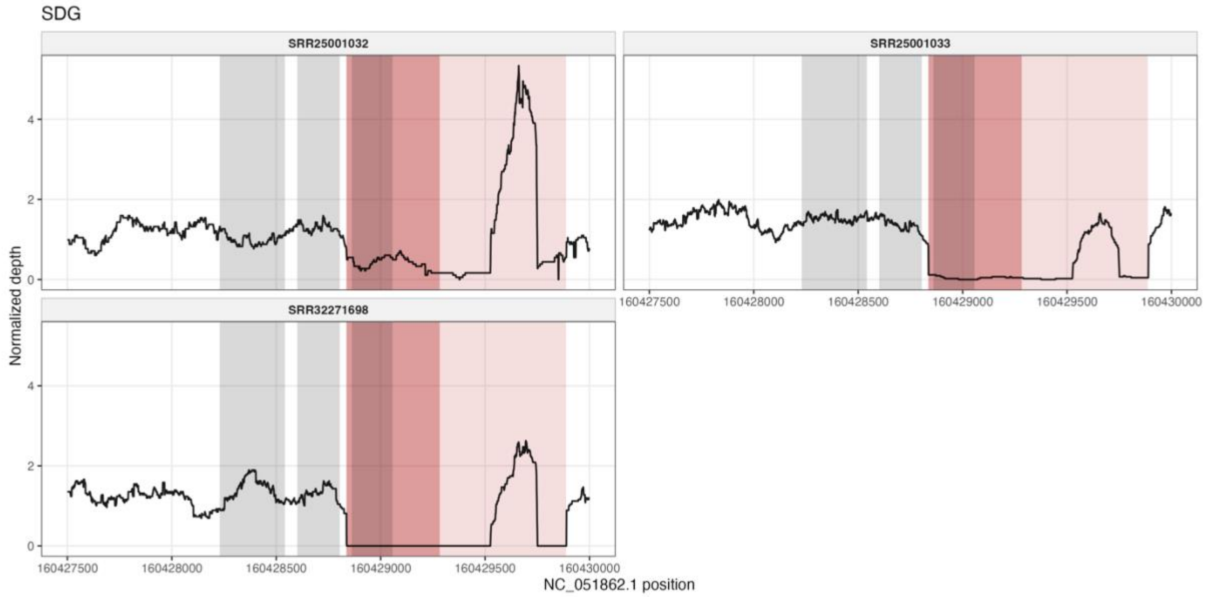


**Supplemental Figure 9:** Normalized read depth plots for the *GST-1* deletion region in each of three individuals from San Diego, California. Read depth was normalized by that individuals average read depth across the entire genome. Data was obtained from Haba et al. 2025. Light red shaded regions indicate the larger deletion while darker shaded region indicates the deletion with an alternative ending and grey shaded regions indicate each exon in the *GST-1* gene.

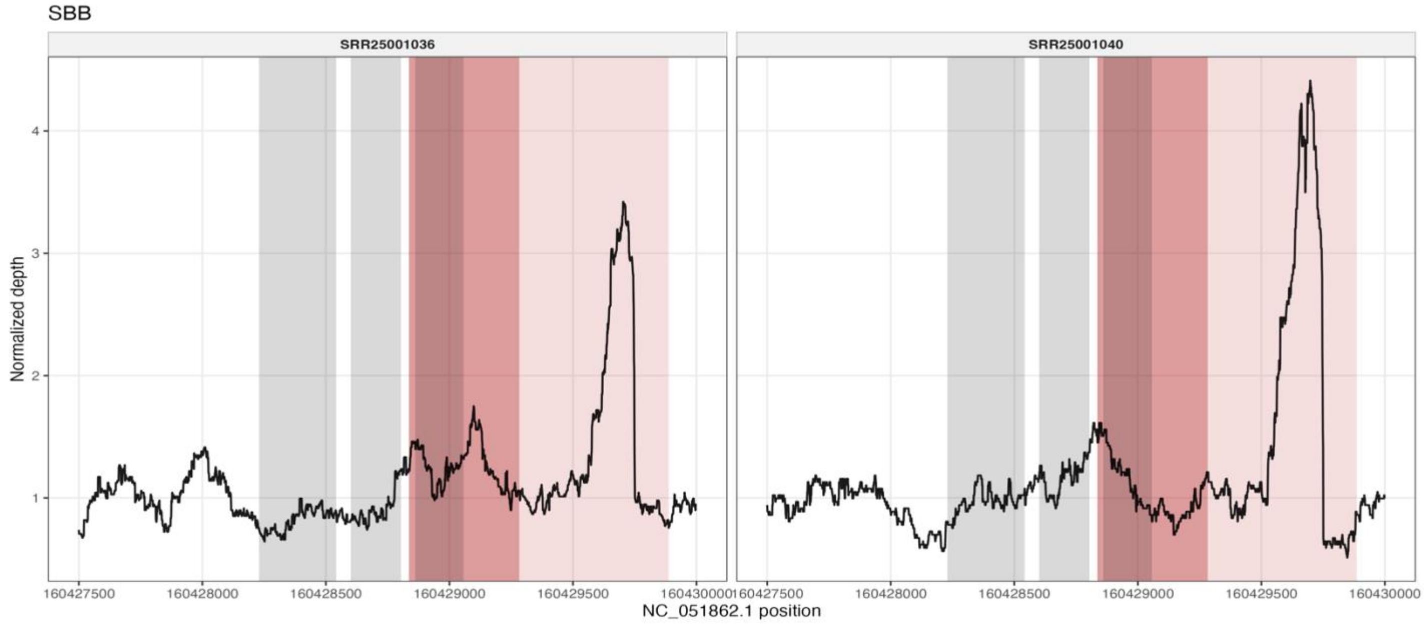


**Supplemental Figure 10:** Normalized read depth plots for the *GST-1* deletion region in each of two individuals from Santa Barbara, California. Read depth was normalized by that individuals average read depth across the entire genome. Data was obtained from Haba et al. 2025. Light red shaded regions indicate the larger deletion while darker shaded region indicates the deletion with an alternative ending and grey shaded regions indicate each exon in the *GST-1* gene.

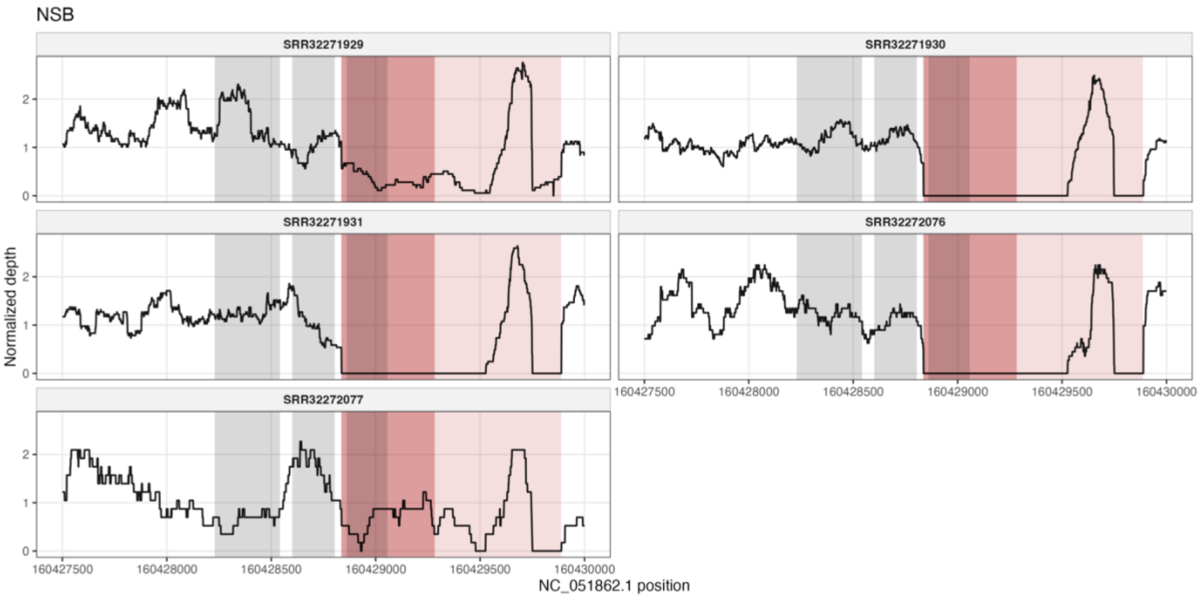


**Supplemental Figure 11:** Normalized read depth plots for the *GST-1* deletion region in each of five individuals from Nosy Be, Madagascar. Read depth was normalized by that individuals average read depth across the entire genome. Data was obtained from Haba et al. 2025. Light red shaded regions indicate the larger deletion while darker shaded region indicates the deletion with an alternative ending and grey shaded regions indicate each exon in the *GST-1* gene.

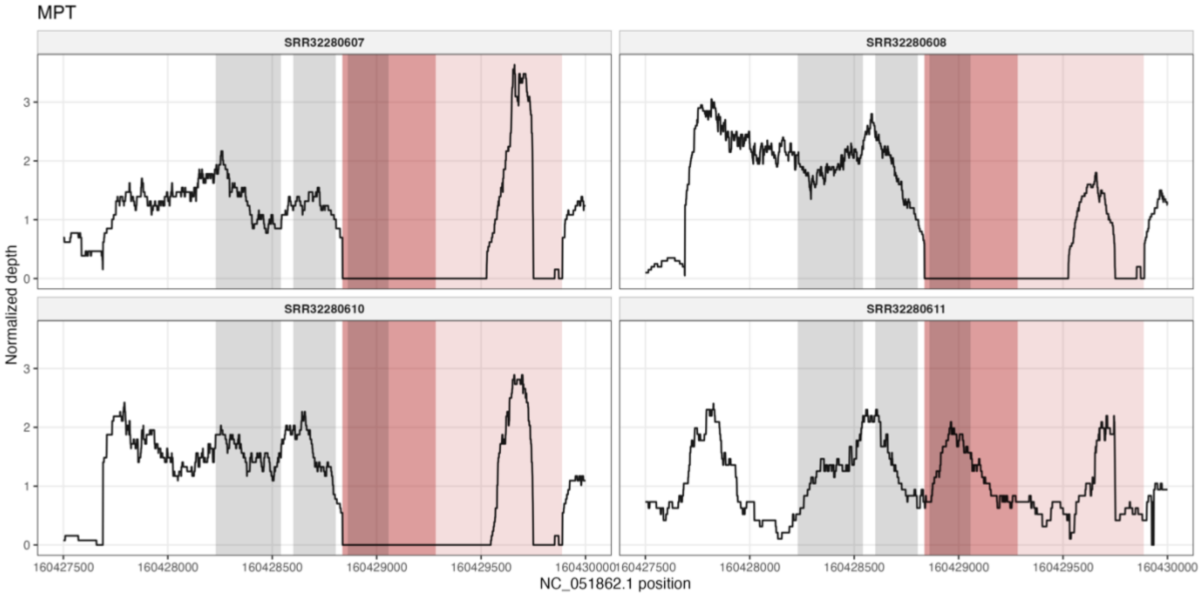


**Supplemental Figure 12:** Normalized read depth plots for the *GST-1* deletion region in each of four individuals from Maputo, Mozambique. Read depth was normalized by that individuals average read depth across the entire genome. Data was obtained from Haba et al. 2025. Light red shaded regions indicate the larger deletion while darker shaded region indicates the deletion with an alternative ending and grey shaded regions indicate each exon in the *GST-1* gene.

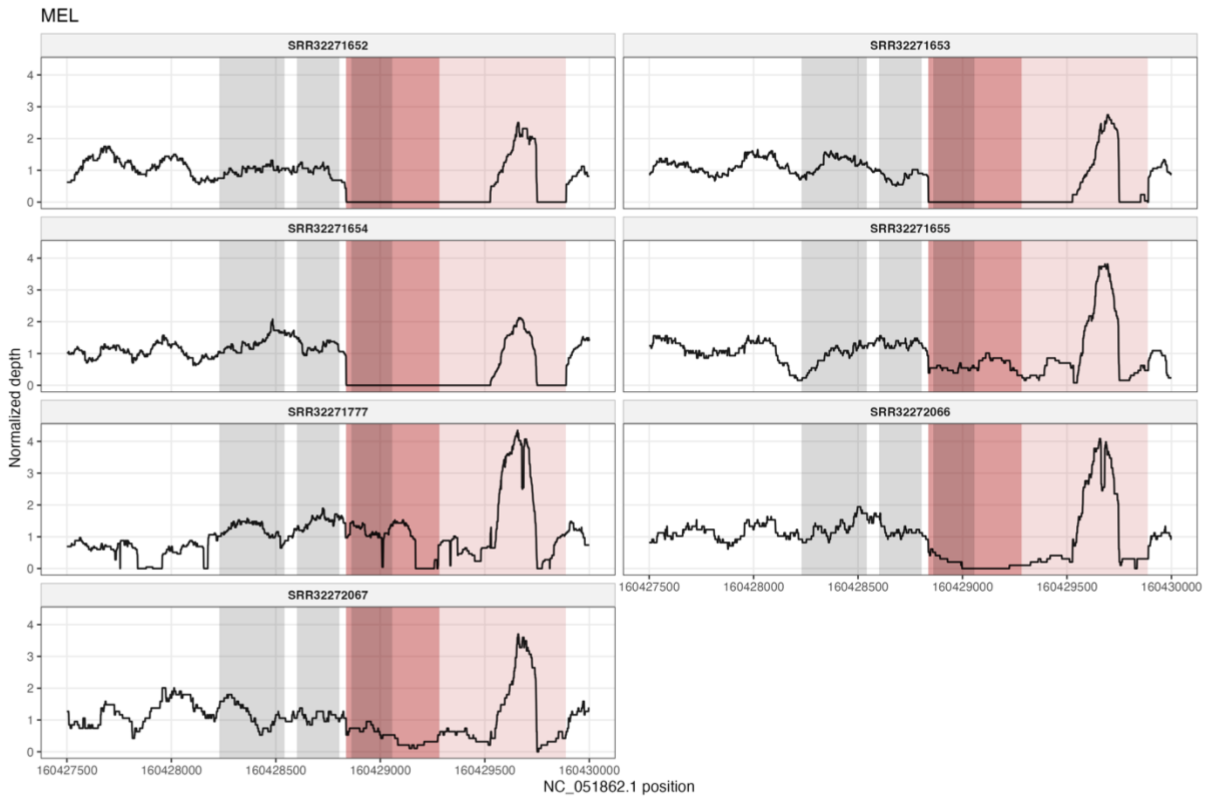


**Supplemental Figure 13:** Normalized read depth plots for the *GST-1* deletion region in each of seven individuals from Melaka, Malaysia. Read depth was normalized by that individuals average read depth across the entire genome. Data was obtained from Haba et al. 2025. Light red shaded regions indicate the larger deletion while darker shaded region indicates the deletion with an alternative ending and grey shaded regions indicate each exon in the *GST-1* gene.

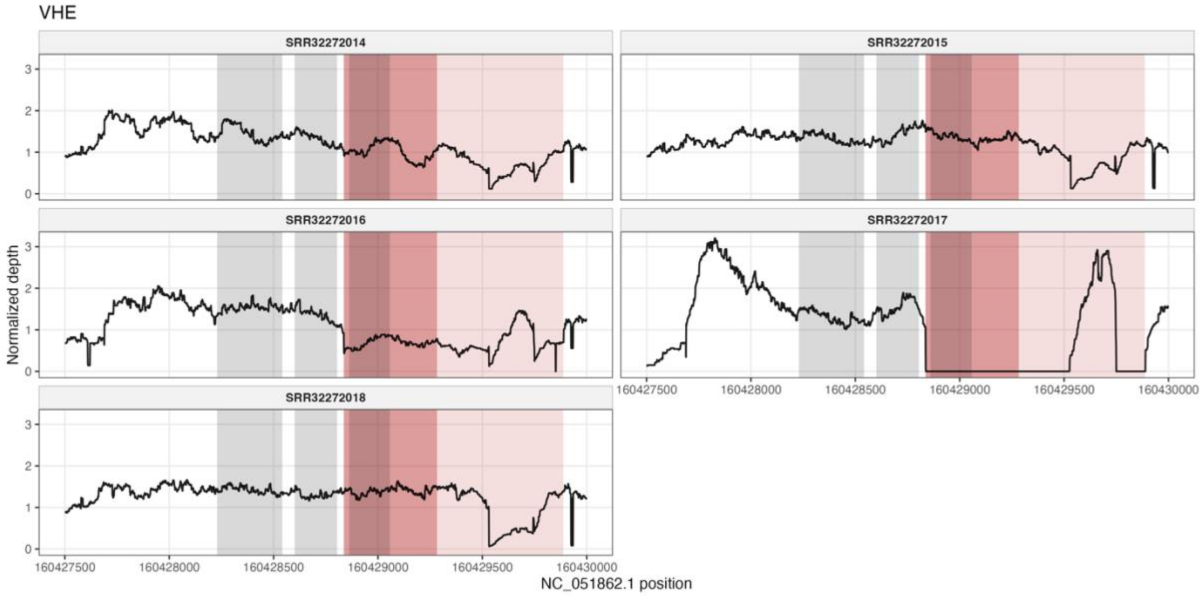


**Supplemental Figure 14:** Normalized read depth plots for the *GST-1* deletion region in each of five individuals from Vhembe, South Africa. Read depth was normalized by that individuals average read depth across the entire genome. Data was obtained from Haba et al. 2025. Light red shaded regions indicate the larger deletion while darker shaded region indicates the deletion with an alternative ending and grey shaded regions indicate each exon in the *GST-1* gene.

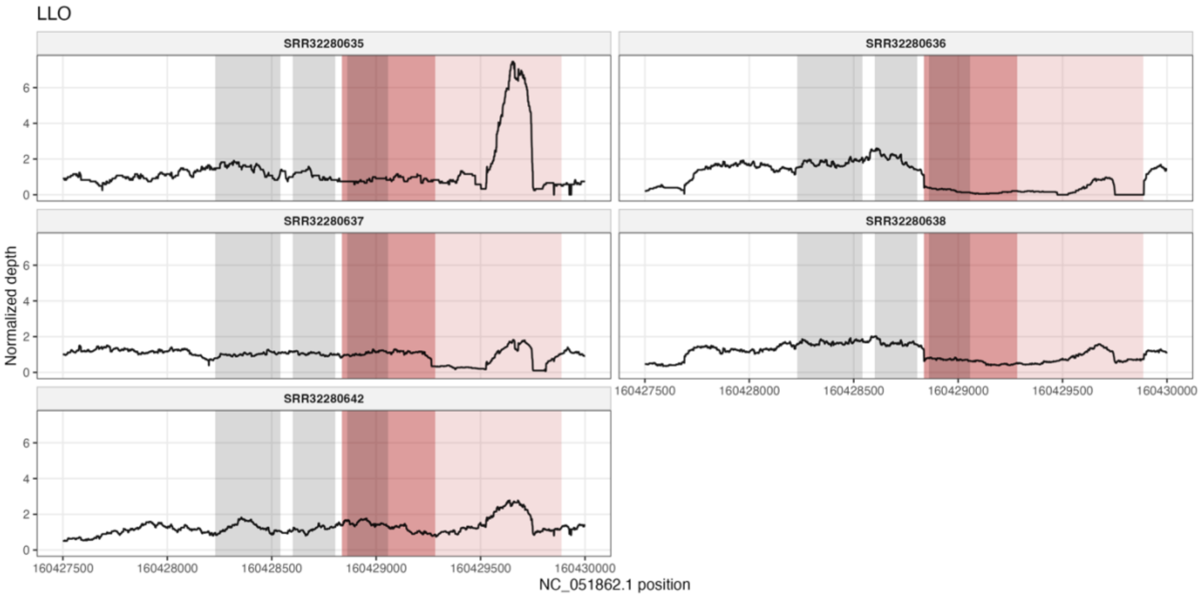


**Supplemental Figure 15:** Normalized read depth plots for the *GST-1* deletion region in each of five individuals from La Lope, Gabon. Read depth was normalized by that individuals average read depth across the entire genome. Data was obtained from Haba et al. 2025. Light red shaded regions indicate the larger deletion while darker shaded region indicates the deletion with an alternative ending and grey shaded regions indicate each exon in the *GST-1* gene.

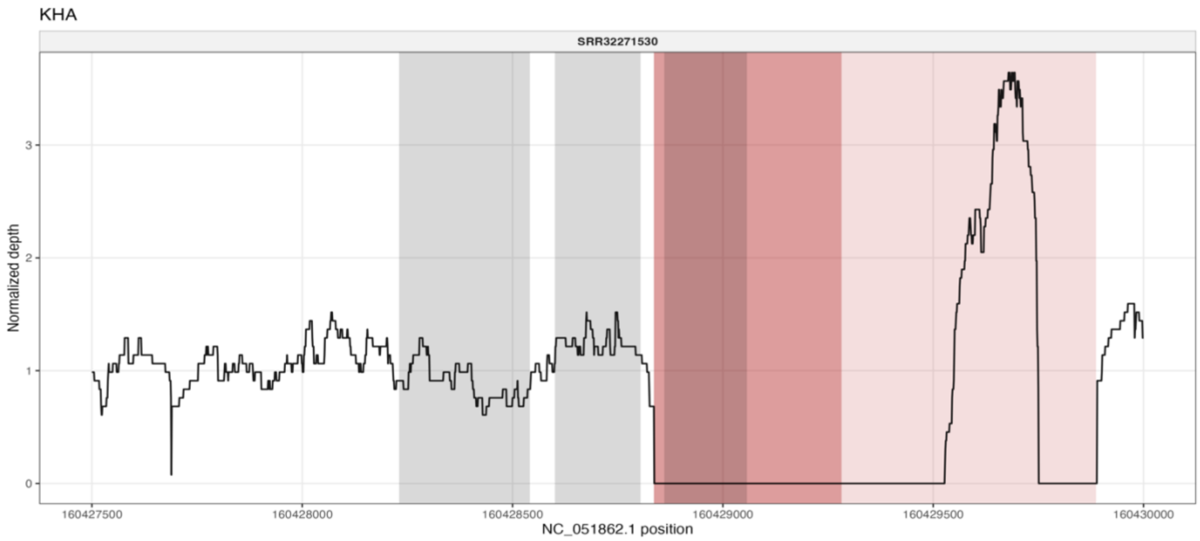


**Supplemental Figure 16:** Normalized read depth plot for the *GST-1* deletion region in one individual from Khartoum, Sudan. Read depth was normalized by that individuals average read depth across the entire genome. Data was obtained from Haba et al. 2025. Light red shaded regions indicate the larger deletion while darker shaded region indicates the deletion with an alternative ending and grey shaded regions indicate each exon in the *GST-1* gene.

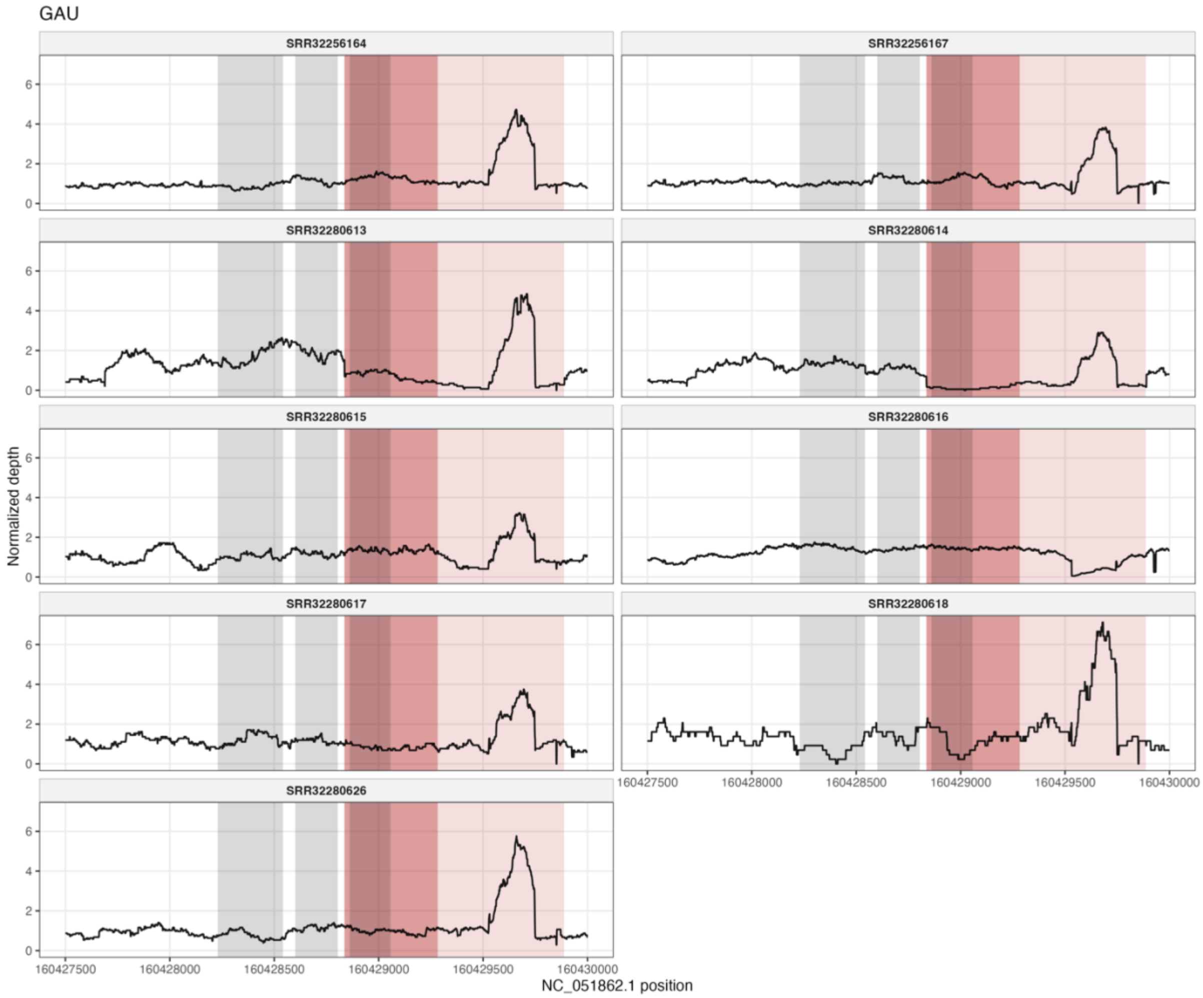


**Supplemental Figure 17:** Normalized read depth plots for the *GST-1* deletion region in each of nine individuals from Gauteng, South Africa. Read depth was normalized by that individuals average read depth across the entire genome. Data was obtained from Haba et al. 2025. Light red shaded regions indicate the larger deletion while darker shaded region indicates the deletion with an alternative ending and grey shaded regions indicate each exon in the *GST-1* gene.

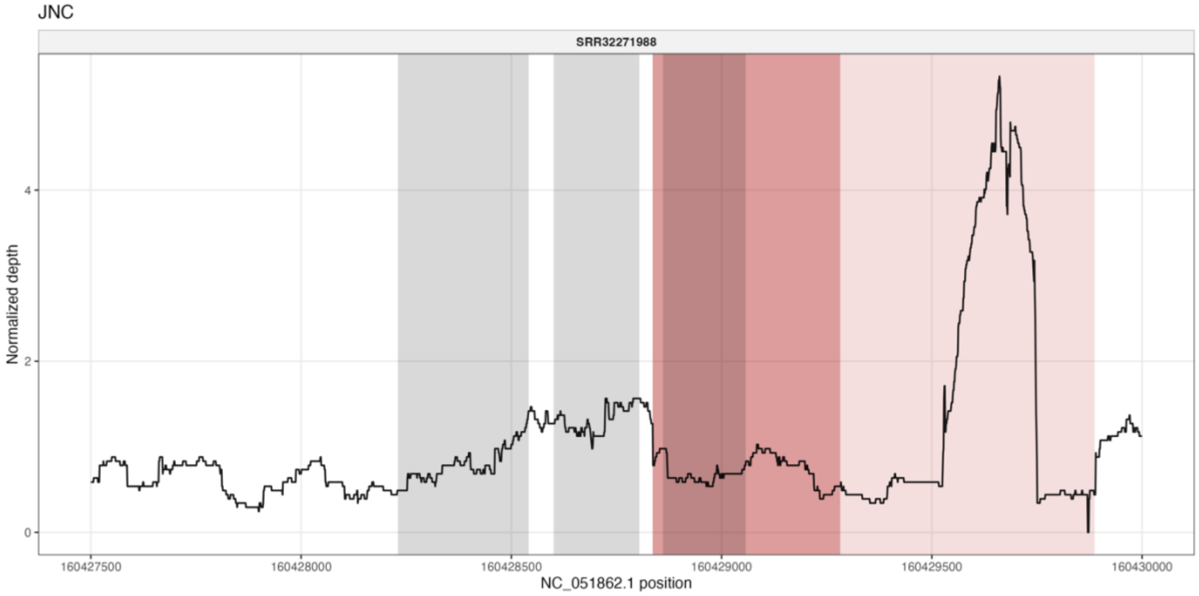


**Supplemental Figure 18:** Normalized read depth plot for the *GST-1* deletion region in one individual from Junin City, Argentina. Read depth was normalized by that individuals average read depth across the entire genome. Data was obtained from Haba et al. 2025. Light red shaded regions indicate the larger deletion while darker shaded region indicates the deletion with an alternative ending and grey shaded regions indicate each exon in the *GST-1* gene.

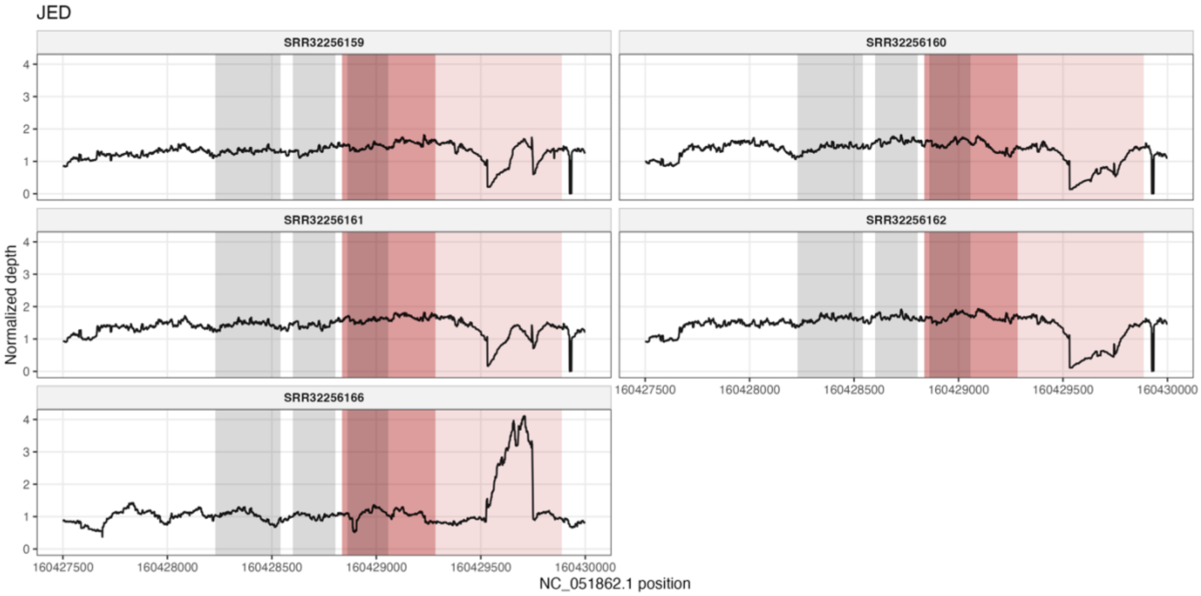


**Supplemental Figure 19:** Normalized read depth plots for the *GST-1* deletion region in each of five individuals from Jeddah, Saudi Arabia. Read depth was normalized by that individuals average read depth across the entire genome. Data was obtained from Haba et al. 2025. Light red shaded regions indicate the larger deletion while darker shaded region indicates the deletion with an alternative ending and grey shaded regions indicate each exon in the *GST-1* gene.

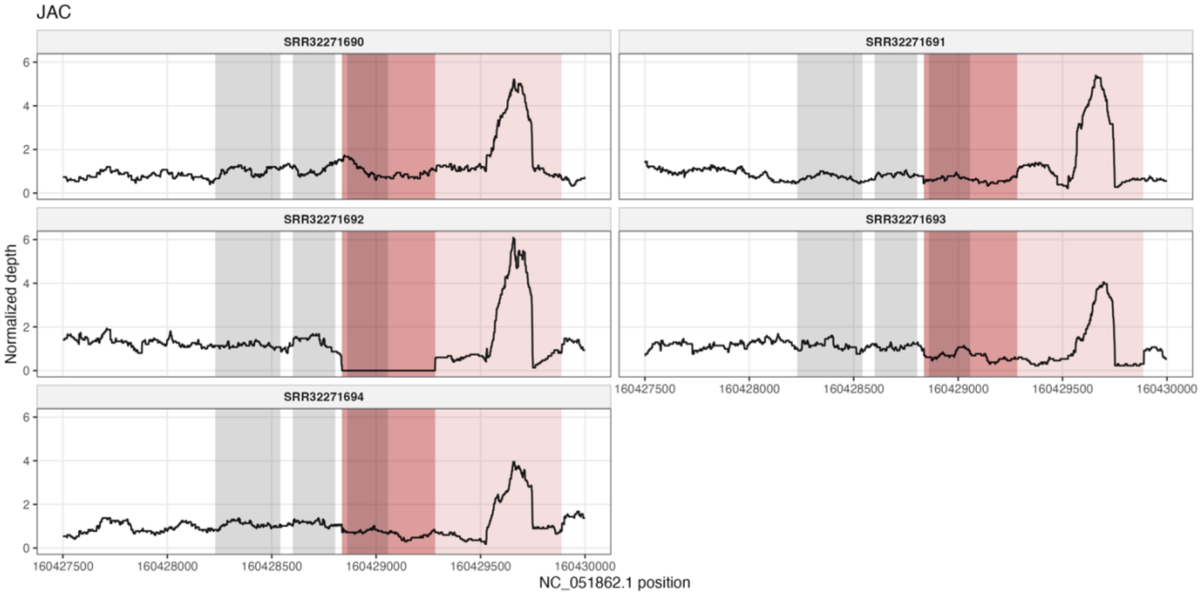


**Supplemental Figure 20:** Normalized read depth plots for the *GST-1* deletion region in each of five individuals from St. Augustine, Florida. Read depth was normalized by that individuals average read depth across the entire genome. Data was obtained from Haba et al. 2025. Light red shaded regions indicate the larger deletion while darker shaded region indicates the deletion with an alternative ending and grey shaded regions indicate each exon in the *GST-1* gene.

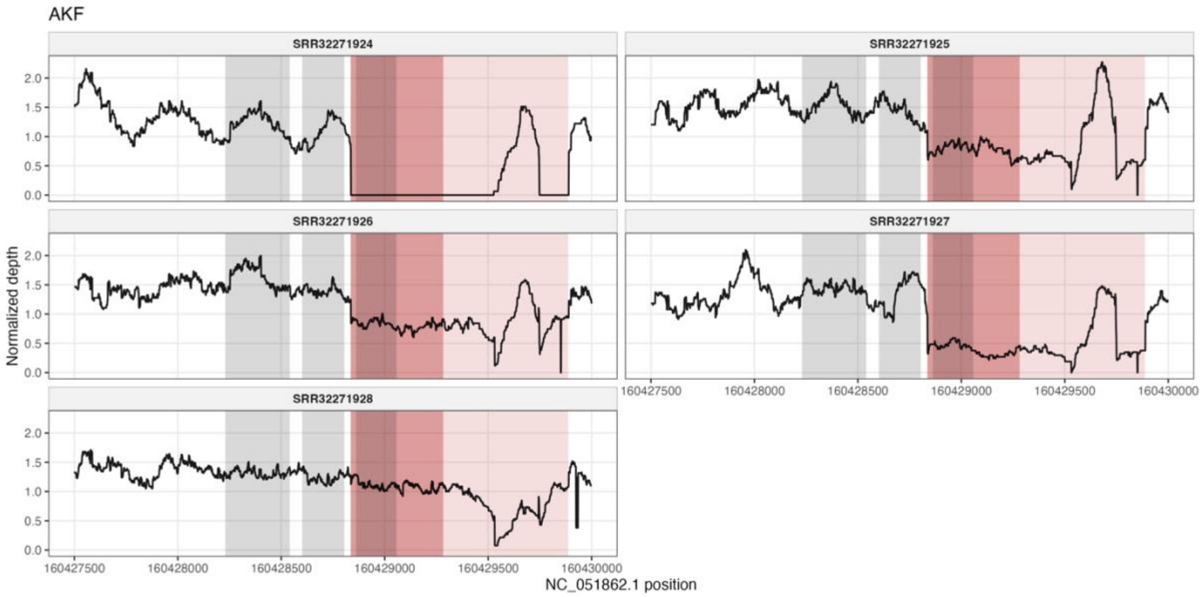


**Supplemental Figure 21:** Normalized read depth plots for the *GST-1* deletion region in each of five individuals from Boeny, Madagascar. Read depth was normalized by that individuals average read depth across the entire genome. Data was obtained from Haba et al. 2025. Light red shaded regions indicate the larger deletion while darker shaded region indicates the deletion with an alternative ending and grey shaded regions indicate each exon in the *GST-1* gene.

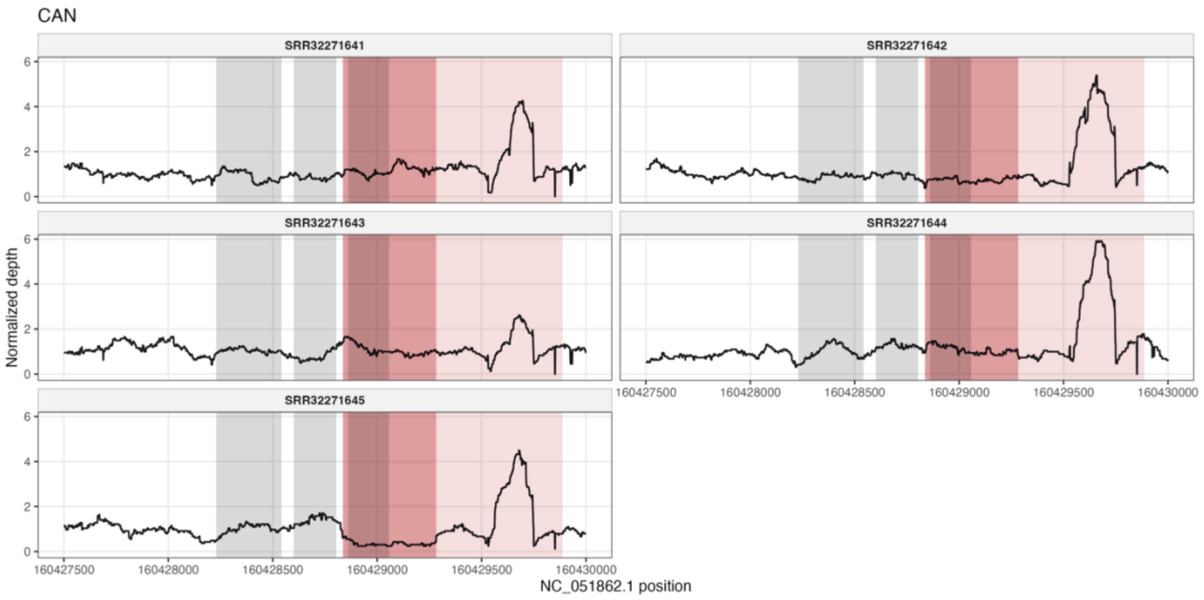


**Supplemental Figure 22:** Normalized read depth plots for the *GST-1* deletion region in each of five individuals from Australian Capital Territory, Australia. Read depth was normalized by that individuals average read depth across the entire genome. Data was obtained from Haba et al. 2025. Light red shaded regions indicate the larger deletion while darker shaded region indicates the deletion with an alternative ending and grey shaded regions indicate each exon in the *GST-1* gene.

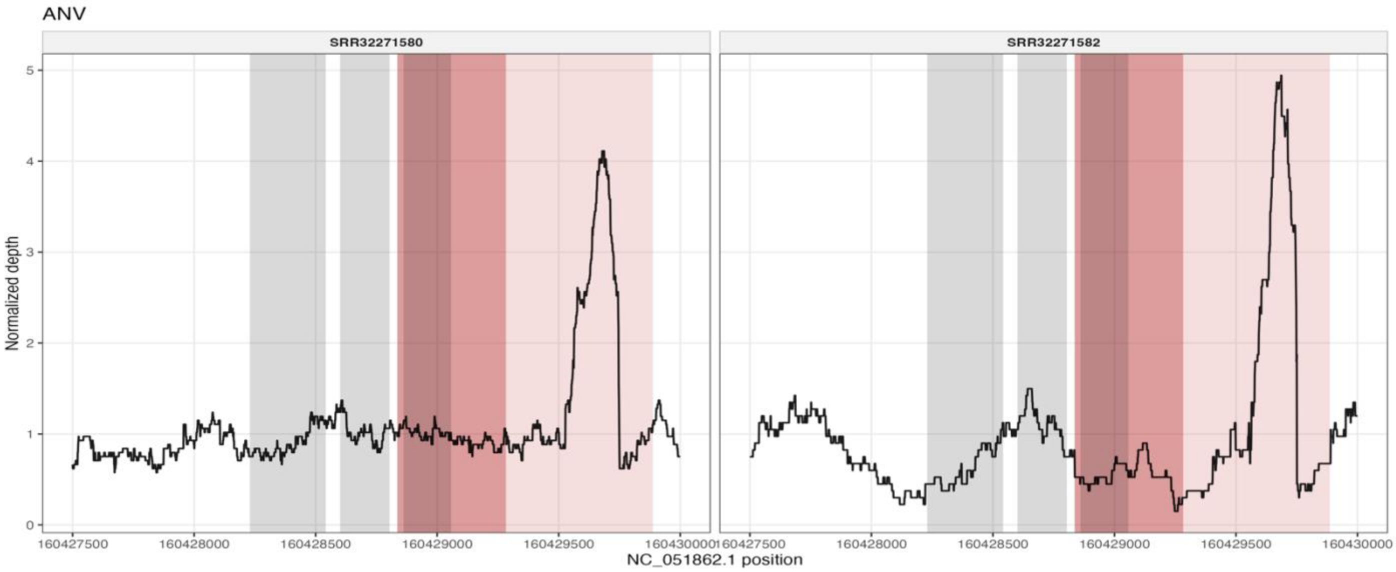


**Supplemental Figure 23:** Normalized read depth plots for the *GST-1* deletion region in each of two individuals from Palmdale, California. Read depth was normalized by that individuals average read depth across the entire genome. Data was obtained from Haba et al. 2025. Light red shaded regions indicate the larger deletion while darker shaded region indicates the deletion with an alternative ending and grey shaded regions indicate each exon in the *GST-1* gene.

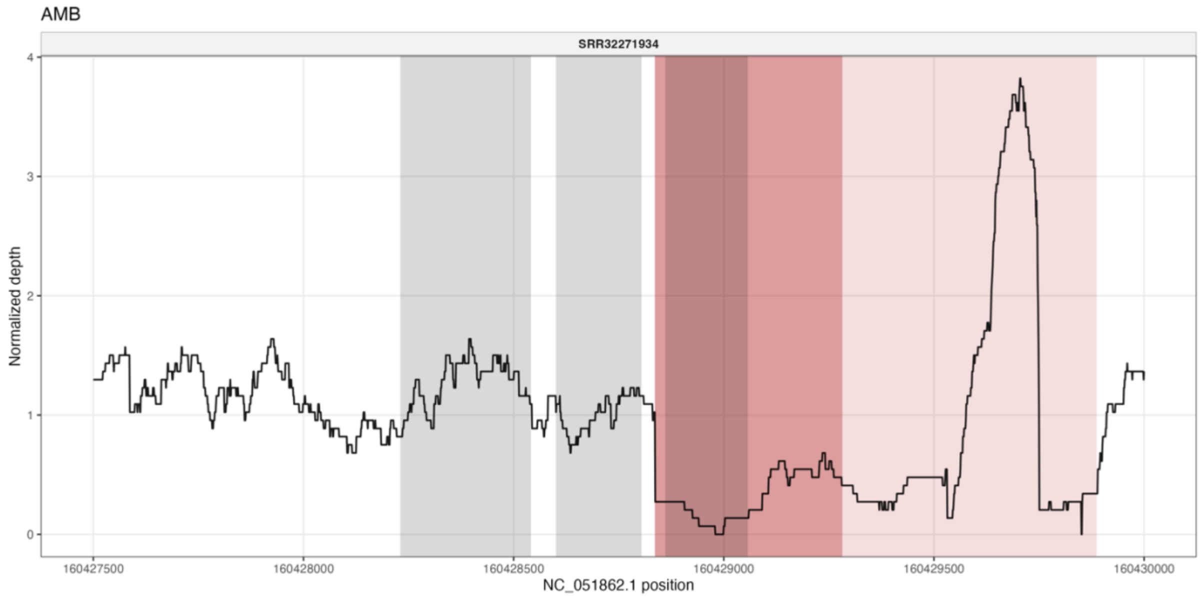


**Supplemental Figure 24:** Normalized read depth plot for the *GST-1* deletion region from one individual Ambalavao, Madagascar. Read depth was normalized by that individuals average read depth across the entire genome. Data was obtained from Haba et al. 2025. Light red shaded regions indicate the larger deletion while darker shaded region indicates the deletion with an alternative ending and grey shaded regions indicate each exon in the *GST-1* gene.

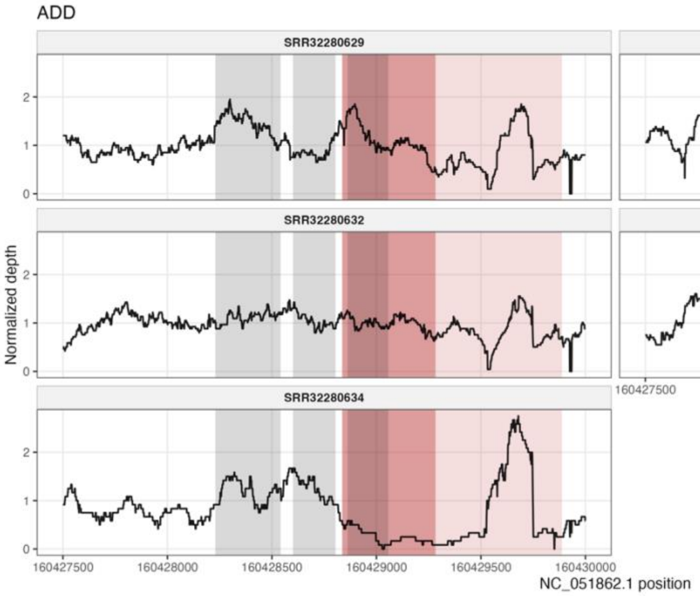


**Supplemental Figure 25:** Normalized read depth plots for the *GST-1* deletion region in each of five individuals from Addis Ababa, Ethiopia. Read depth was normalized by that individuals average read depth across the entire genome. Data was obtained from Haba et al. 2025. Light red shaded regions indicate the larger deletion while darker shaded region indicates the deletion with an alternative ending and grey shaded regions indicate each exon in the *GST-1* gene.

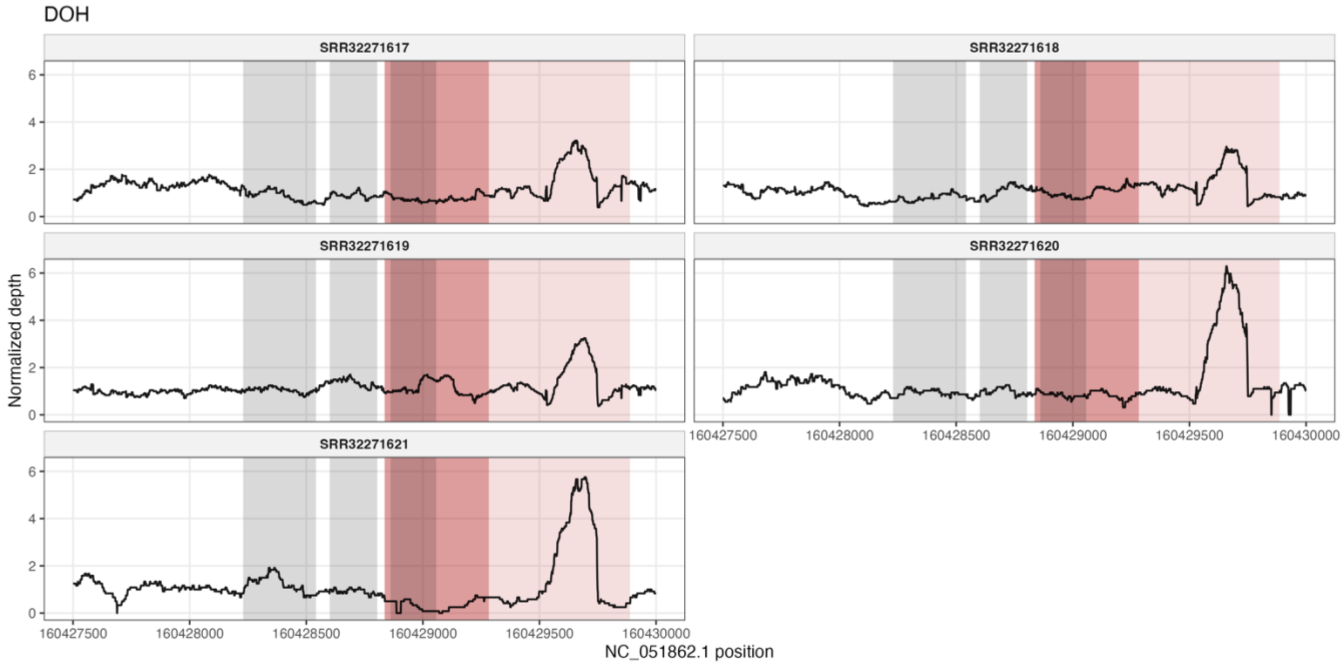


**Supplemental Figure 26:** Normalized read depth plots for the *GST-1* deletion region in each of five individuals from Doha, Qatar. Read depth was normalized by that individuals average read depth across the entire genome. Data was obtained from Haba et al. 2025. Light red shaded regions indicate the larger deletion while darker shaded region indicates the deletion with an alternative ending and grey shaded regions indicate each exon in the *GST-1* gene. Note that we collapsed the samples collected from Doha (SRR32271620-21) in with those collected from Al Rayyan (SRR32271617-19), due to their geographic proximity.

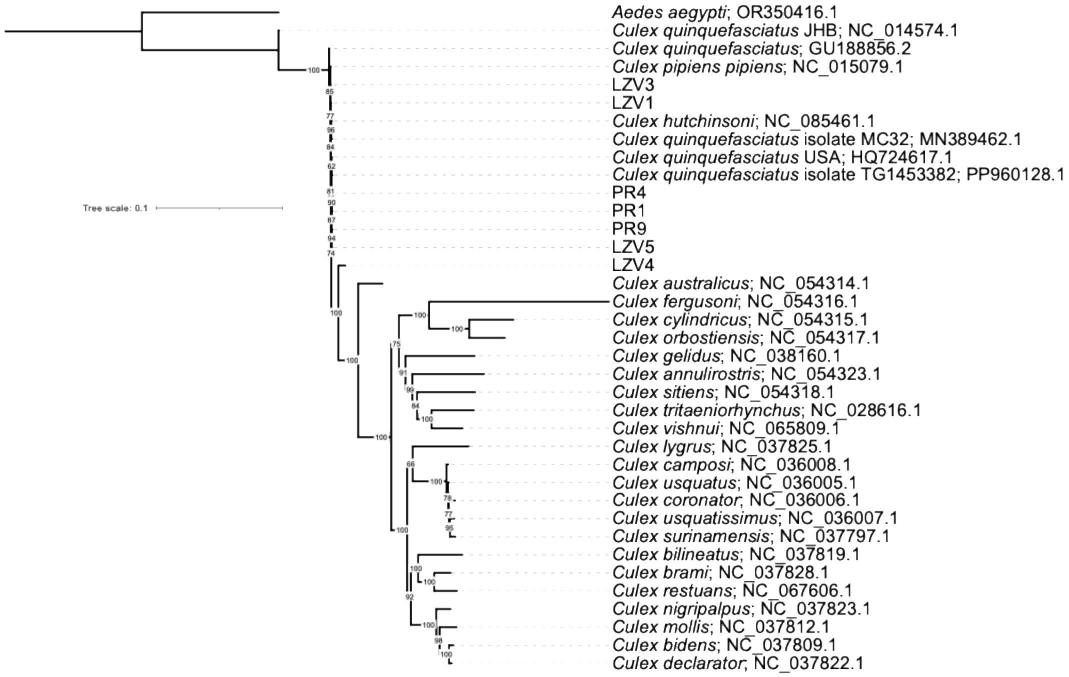


**Supplemental Figure 27.** Phylogeny generated from mitochondrial genomes of the seven *C. quinquefasciatus* samples sequenced for this study, along with 29 published *Culex* mitochondrial genomes and an *Aedes aegypti* mitogenome as the outgroup. This phylogeny was based on an alignment of approximately 16,200 bp.

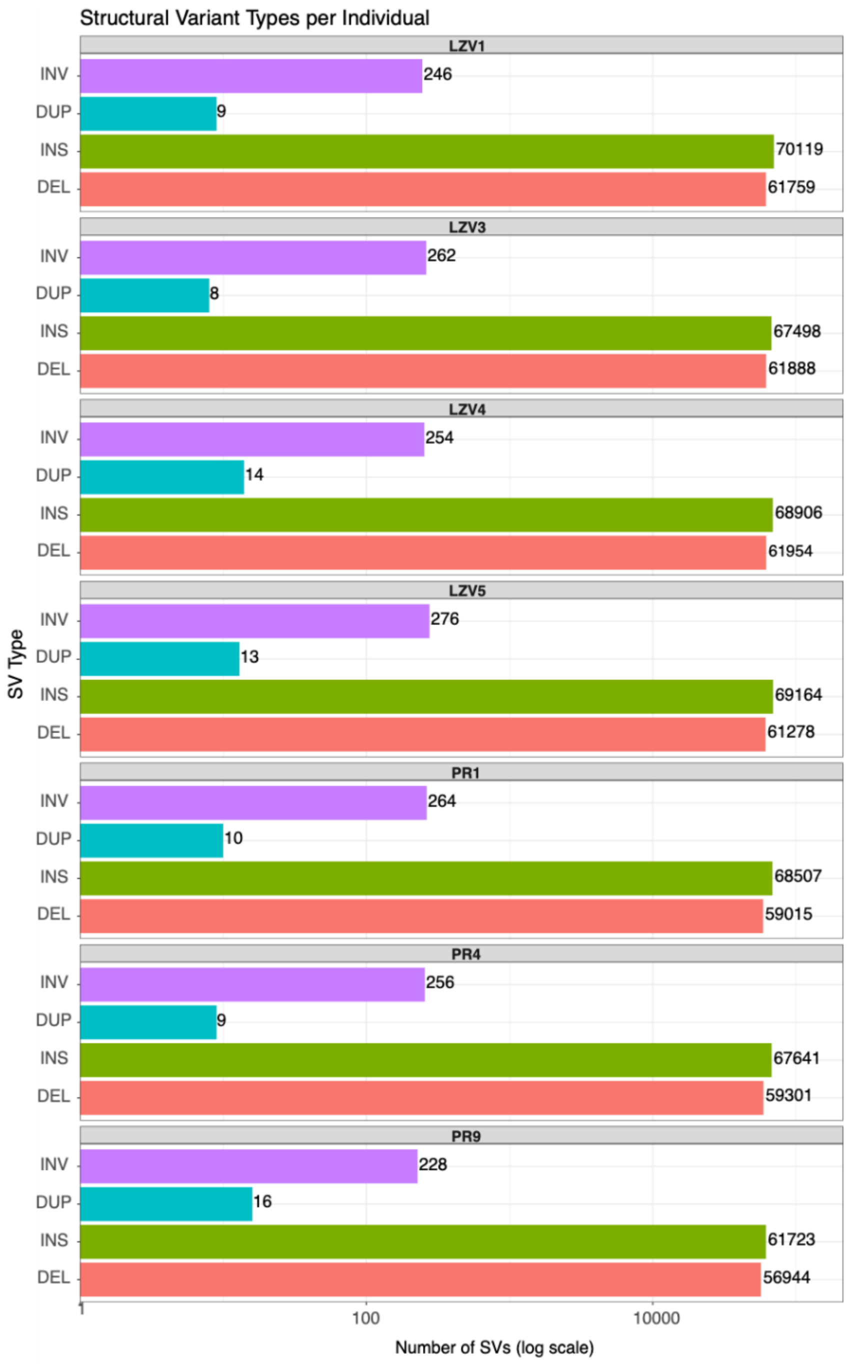


**Supplemental Figure 28.** Counts of structural variant types present in each individual.

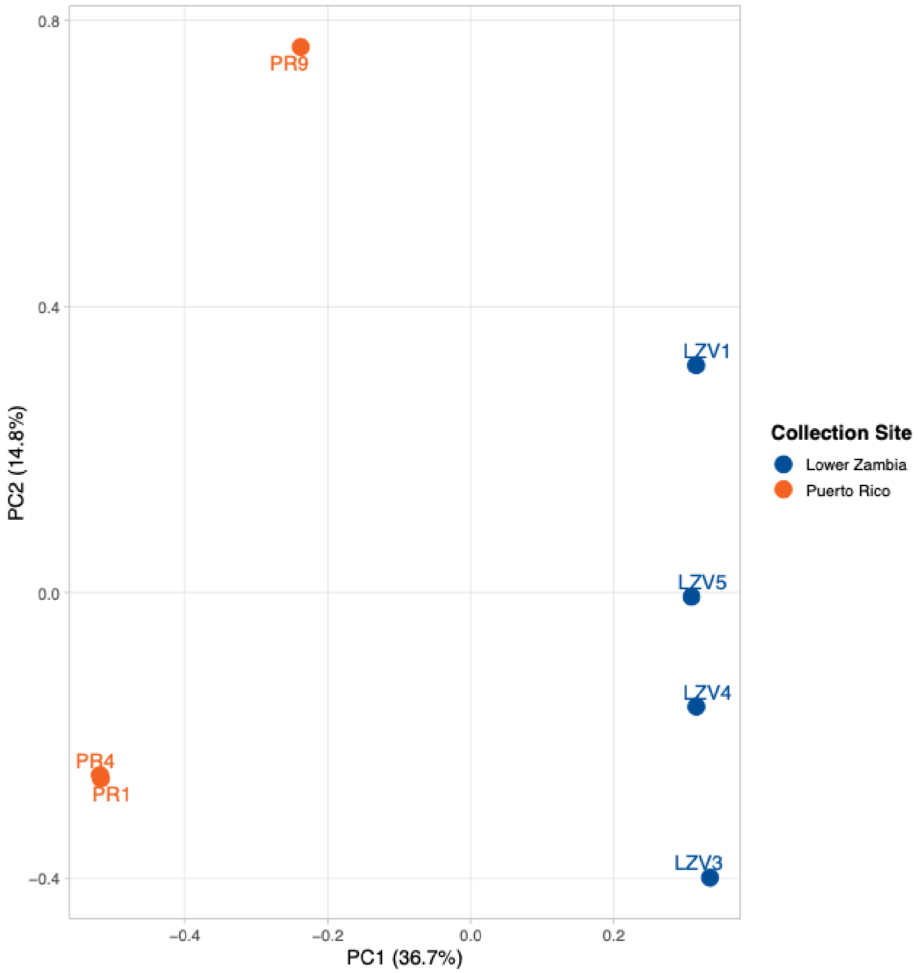


**Supplemental Figure 29.** Principal Component Analysis (PCA) of SNPs showing population structure of Zambian and Puerto Rican samples.

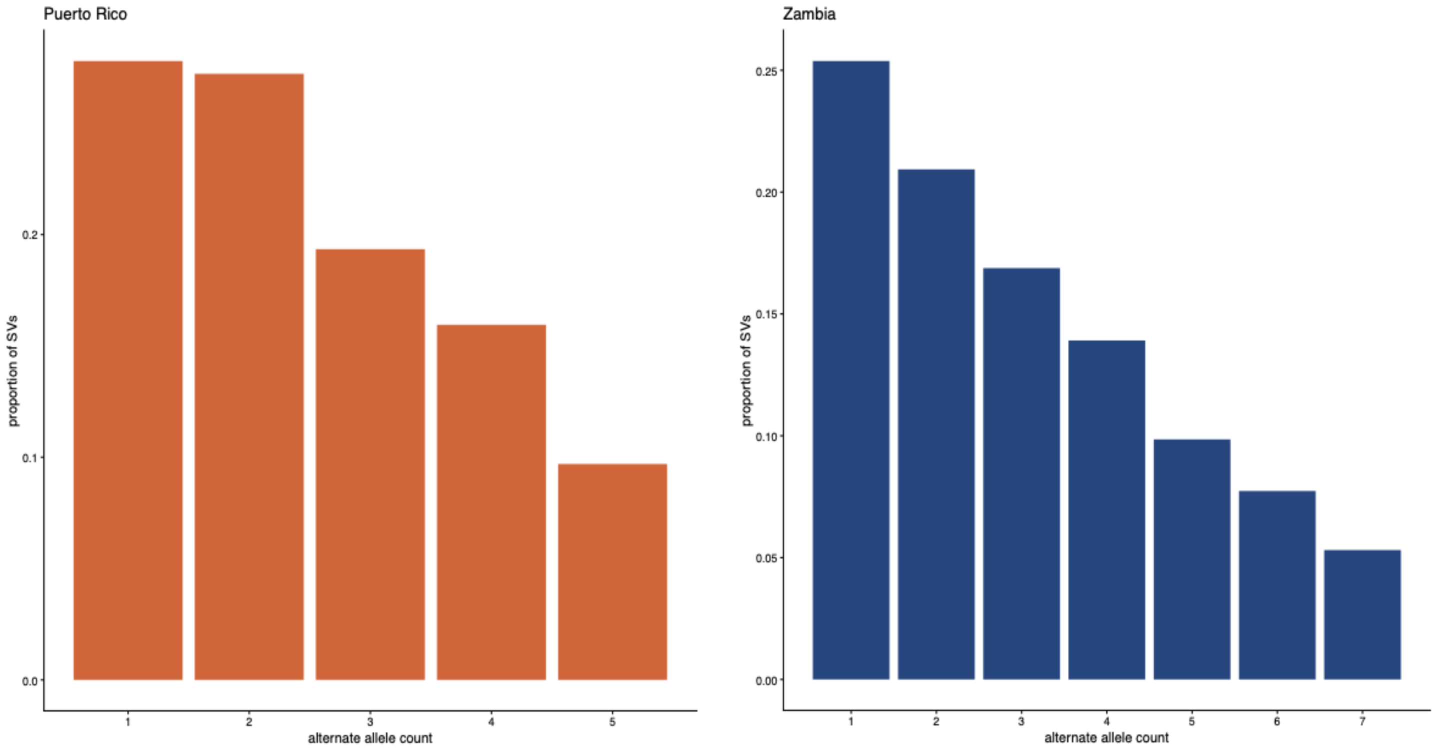


**Supplemental Figure 30.** Site frequency spectra (SFS) generated from the full set of SVs with missing and fixed genotypes removed for Puerto Rico (left) and Zambia (right).

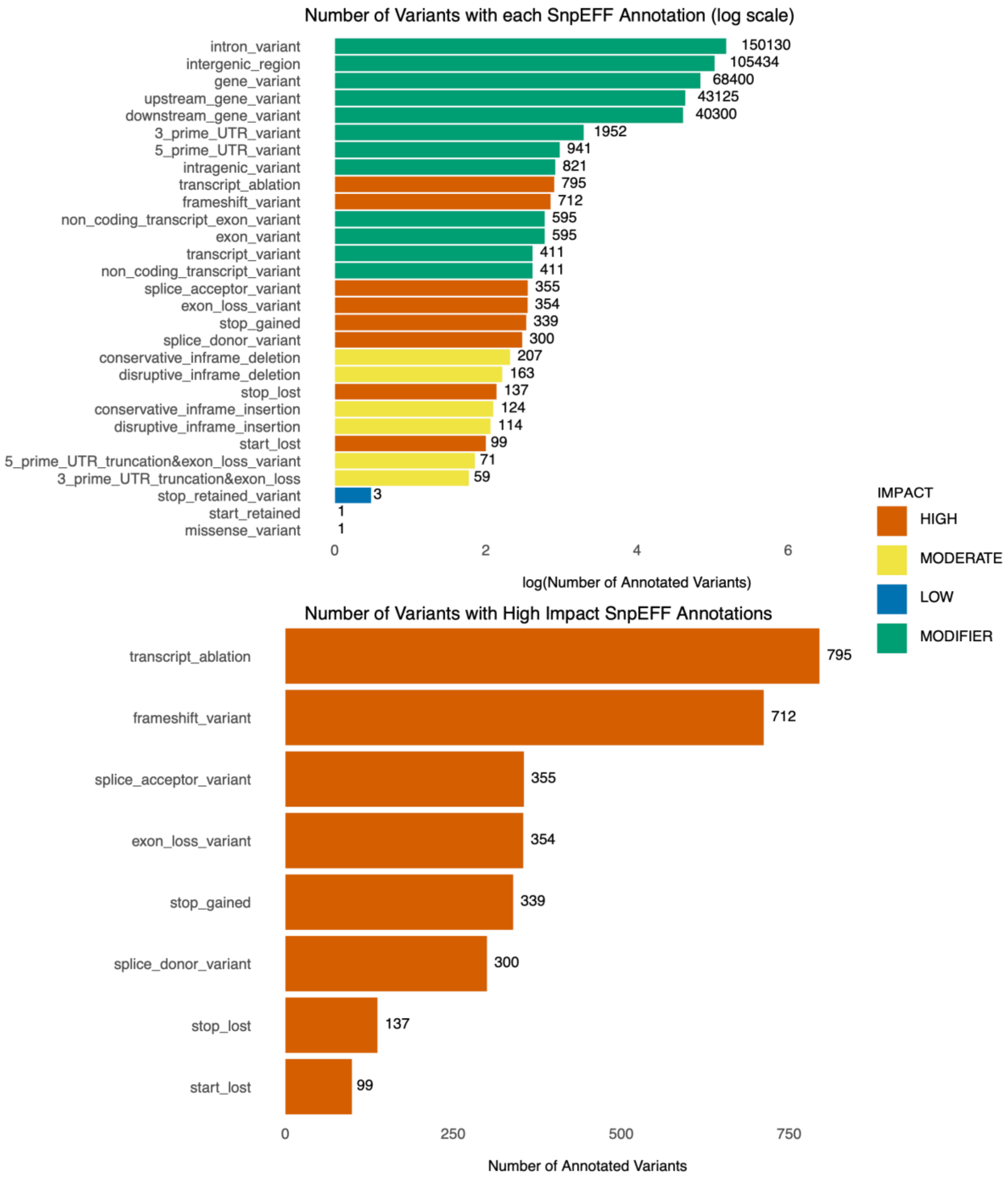


**Supplemental Figure 31.** (Top) The number of SVs grouped by SnpEff annotation and the number of SVs grouped by SnpEff annotation labelled as 'high impact' variants (bottom).

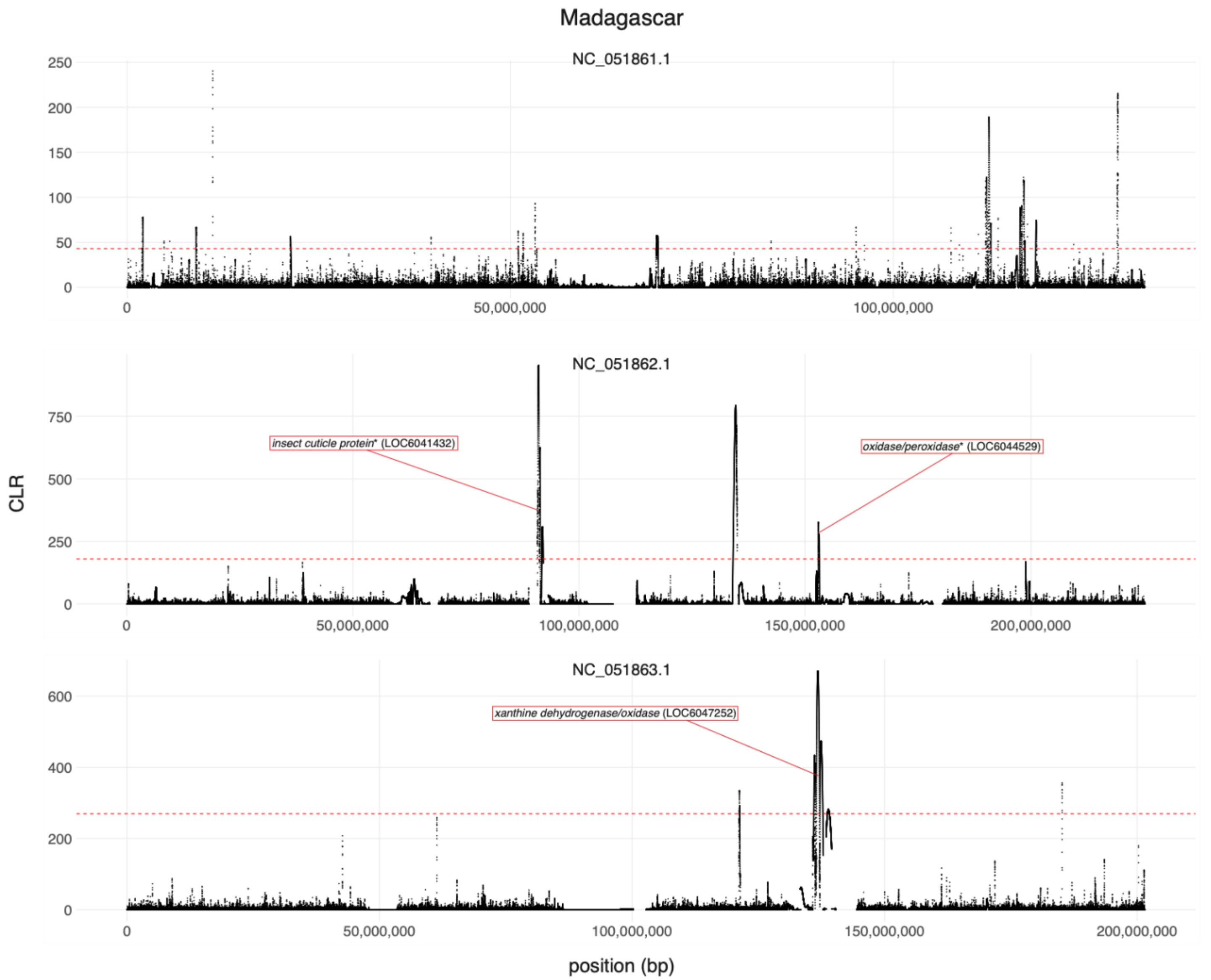


**Supplemental Figure 32.** Insecticide resistance candidate genes overlapping both CLR peaks and SVs identified by Sweepfinder2 in *Cx. quinquefasciatus* populations from Madagascar. Pictured genes overlap SVs that are present in at least one individual from the Zambian population. SweepFinder analysis was performed using sequencing data from Haba et al (2025). The dashed red line indicates the 99$^{th}$ percentile of CLR values across the chromosome, which was used to define a CLR peak. Genes with asterisks were annotated as uncharacterized loci by NCBI, but we included annotation information from the conserved domain database here (Wang et al. 2023).

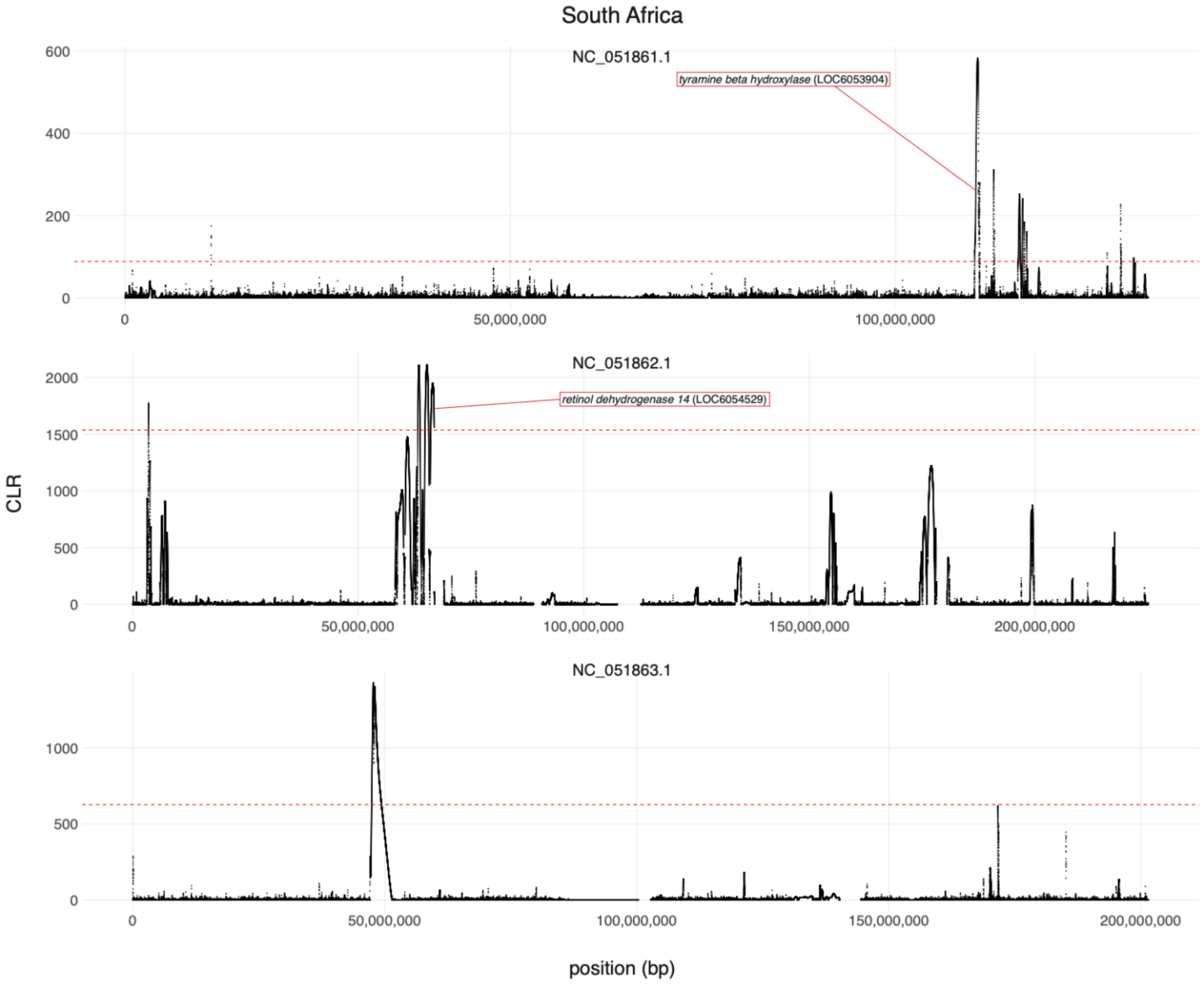


**Supplemental Figure 33.** Insecticide resistance candidate genes overlapping both CLR peaks and SVs identified by Sweepfinder2 in *Cx. quinquefasciatus* populations from South Africa. Pictured genes overlap SVs that are present in at least one individual from the Zambian population. SweepFinder analysis was performed using sequencing data from Haba et al (2025). The dashed red line indicates the $99^{th}$ percentile of CLR values across the chromosome, which was used to define a CLR peak.

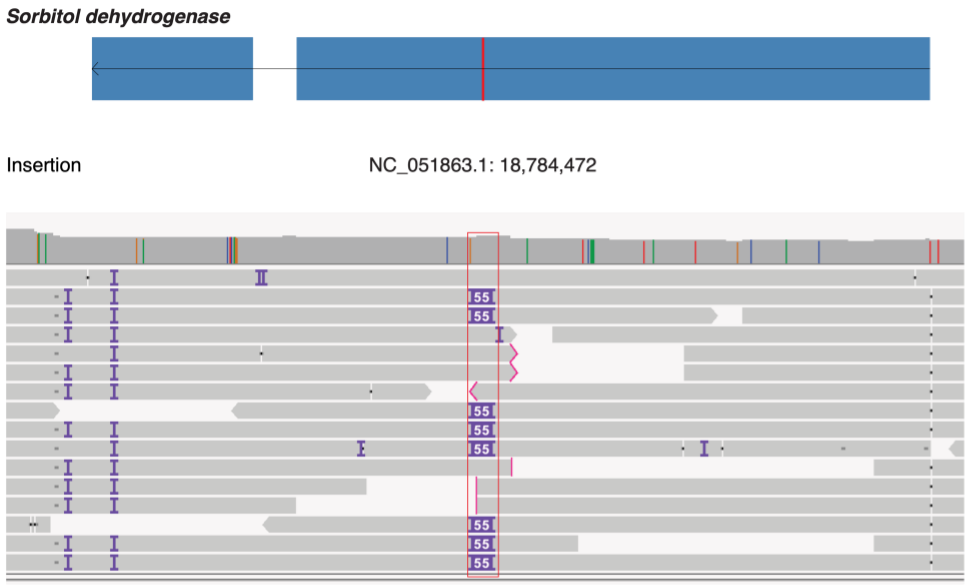


| sample | LZV1 | LZV3 | LZV4 | LZV5 | PR1 | PR4 | PR9 |
|---|---|---|---|---|---|---|---|
| genotype | 0/1 | 0/1 | 0/1 | 1/1 | 0/0 | 0/0 | 0/0 |

**Supplemental Figure 34.** IGV track shown for a structural variant location that was predicted to have high functional impact by SnpEff and overlapped an insecticide resistance gene, *sorbitol dehydrogenase*. The location of the insertion is shown with the red box and the genotypes are recorded at the bottom.

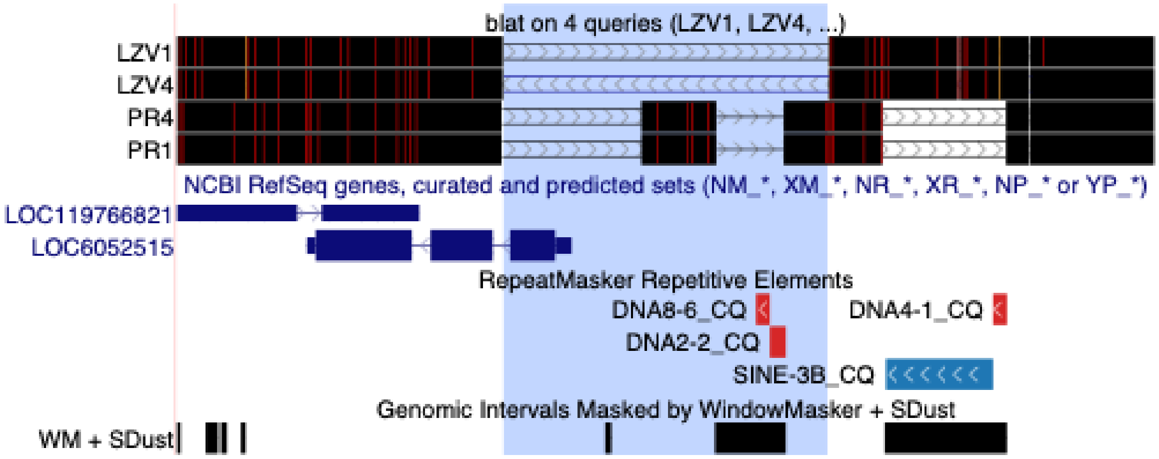


**Supplemental Figure 35.** UCSC Genome Browser shot showing alignments of reads (black bars at the top) from four different genomes: two from Zambia (LZV1 and LZV4) that have the deletion according to our pipeline, and two from Puerto Rico (PR1 and PR4) that do not. The coordinates of the deletion are shaded in blue and correspond perfectly with a gap in the alignments of the LZV genomes' reads, providing strong support for the deletion. This 1,053 bp deletion encompasses the entirety of the first coding exon of GST1 (LOC6052515), probably resulting in complete gene inactivation in homozygotes. PR1 and PR4's reads do not support the presence of this deletion, but instead suggest the presence another complex SV which also appears to delete the first exon of the gene, while inserting additional sequence nearby (not shown). PR1 and PR4 also appear to lack the SINE element shown in blue.

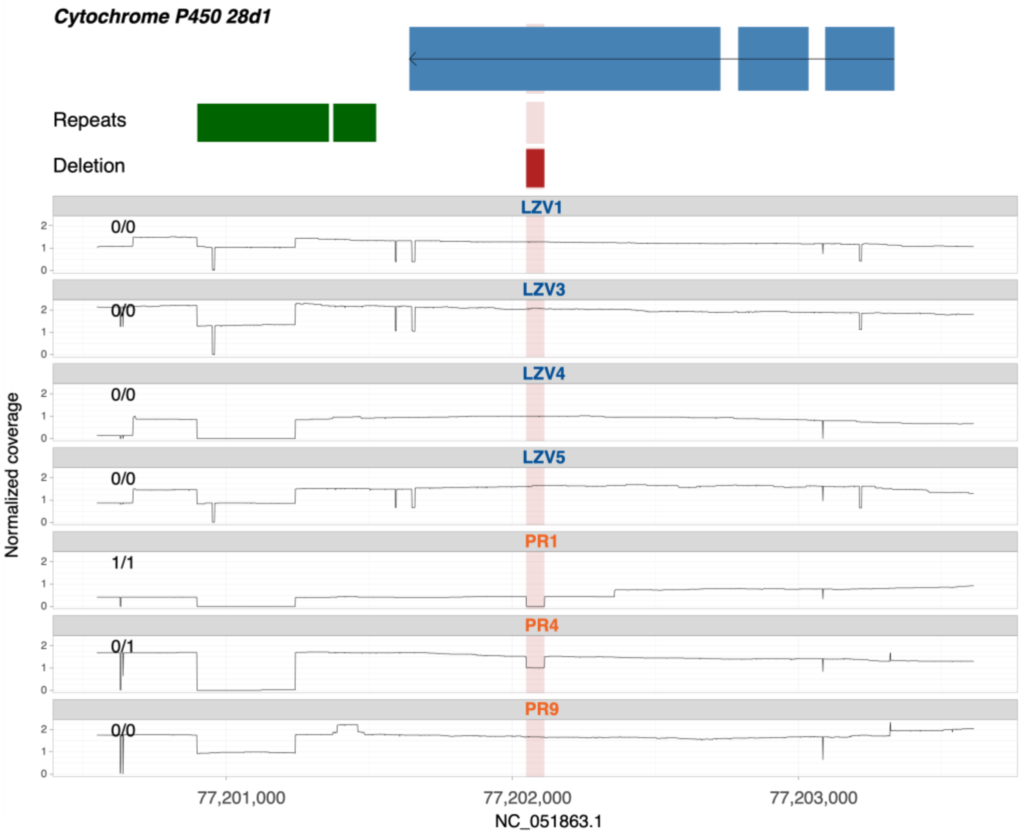


**Supplemental Figure 36.** Normalized read depth plots for all seven individuals at a structural variant location that was predicted to have a high functional impact by SnpEff and overlapped an insecticide resistance gene, *Cytochrome P450 28d1*. Genotypes are recorded to the top left of the read depth plots, gene location and strand direction are in blue at the top, repeats are annotated in green, and the deletion is shown in red.

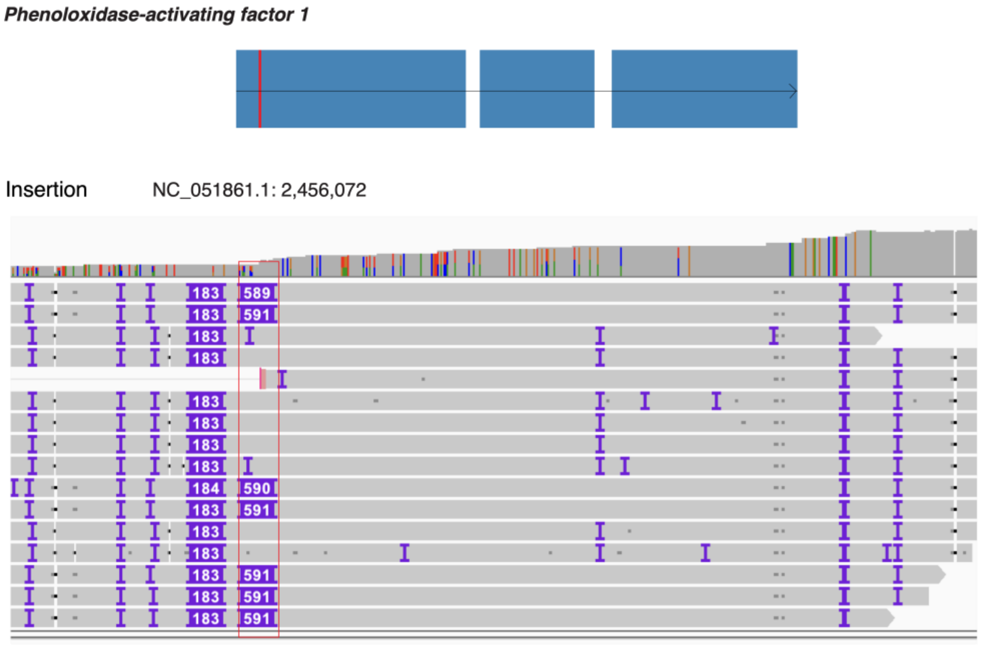


| sample | LZV1 | LZV3 | LZV4 | LZV5 | PR1 | PR4 | PR9 |
|---|---|---|---|---|---|---|---|
| genotype | 0/1 | 0/1 | 0/1 | 1/1 | 0/0 | 0/0 | 0/0 |

**Supplemental Figure 37.** An insertion that overlaps an environmental sensing gene and was predicted to have high functional impact by SnpEff. The IGV track is shown with the insertion highlighted with a red box and the genotypes are recorded at the bottom.

**Supplemental Tables**

**Supplemental Table 1.** Candidate insecticide resistance (IR) gene list consisting of all *Culex quinquefasciatus* genes that correspond to orthologs of putative IR genes in *Anopheles gambiae* published by the *Anopheles Gambiae* 1000 Genomes Consortium at https://malariagen.github.io/agam-selection-atlas/0.1-alpha3/ir-candidates.html. Genes are divided by resistance mechanism: metabolic, target site, behavioral, and cuticular. The table includes gene descriptions, gene IDs, coordinates, strand, as well as corresponding protein IDs. This table also contains the orthogroups output produced by OrthoFinder, which was used to identify *C. quinquefasciatus* genes that are orthologs of previously implicated *An. gambiae* IR genes.

**Supplemental Table 2.** Environmental-related gene list consisting of all *Culex quinquefasciatus* genes that correspond to orthologs from the *Aedes aegypti* chemosensory genes curated in Matthews et al. 2016. We also included odorant receptors, heat shock, ionotropic receptors, and immune genes in our list. The table includes gene descriptions, gene IDs, coordinates, and strand.

**Supplemental Table 3.** Predicted genotypes compared to manually curated genotypes for each of the structural variants highlighted within the manuscript. Table includes the location of the structural variant, the type, size, impacted genes, and the genotypes predicted by the variant callers and the corrected genotypes following manual curation in IGV.

**Supplemental Table 4.** The number of structural variants and the types of variants in each dataset.

| Dataset | Number of SVs | Insertions | Deletions | Duplications | Inversions |
|---|---|---|---|---|---|
| **Full set** | 259,046 | 143,068 | 115,002 | 47 | 929 |
| **High impact** | 2,161 | 426 | 1,710 | 2 | 23 |
| **Sweep** | 2,314 | 1,099 | 1,209 | 1 | 5 |
| **High impact sweep** | 45 | 10 | 35 | 0 | 0 |
| **LZV-enriched** | 13,428 | 6,905 | 6,519 | 0 | 4 |
| **PR-enriched** | 13,899 | 7,596 | 6,299 | 0 | 4 |
| **LZV/PR-enriched** | 27,327 | 14,501 | 12,818 | 0 | 8 |

**Supplemental Table 5.** Genes impacted by structural variants that have predicted high impact effects. Table includes gene ID, structural variant type, transcript ID, gene product, biotype, and gene coordinates.

**Supplemental Table 6.** Number of loss-of-function (LOF) variants predicted by SnpEff for structural variants (SVs) and single nucleotide polymorphisms (SNPs).

| Genome | # of LOF SVs/SNPs | | # of homozygous LOF SVs/SNPs | | # of heterozygous LOF SVs/SNPs | | # of genes affected by SVs/SNPs | | # of genes affected by homozygous LOF SVs/SNPs | |
|---|---|---|---|---|---|---|---|---|---|---|
| | SVs | SNPs | SVs | SNPs | SVs | SNPs | SVs | SNPs | SVs | SNPs |
| LZV1 | 532 | 1110 | 172 | 398 | 360 | 712 | 545 | 798 | 178 | 328 |
| LZV3 | 568 | 1078 | 201 | 414 | 367 | 664 | 578 | 772 | 211 | 349 |
| LZV4 | 562 | 1111 | 180 | 405 | 382 | 706 | 588 | 804 | 202 | 342 |
| LZV5 | 542 | 1109 | 179 | 382 | 363 | 727 | 559 | 819 | 186 | 320 |
| PR1 | 504 | 968 | 193 | 377 | 311 | 591 | 511 | 726 | 201 | 315 |
| PR4 | 502 | 951 | 180 | 403 | 322 | 548 | 514 | 711 | 187 | 336 |
| PR9 | 513 | 986 | 201 | 382 | 312 | 604 | 533 | 706 | 210 | 314 |

**Supplemental Table 7.** Genes impacted by structural variants in the high impact sweep dataset (structural variants with predicted high impact effects that overlap CLR peaks). Table includes gene ID, structural variant type, transcript ID, gene product, biotype, and gene coordinates.

**Supplemental Table 8.** Enrichment of GO terms in structural variants (SV) for each SV dataset, which includes high impact, high impact sweep, LZ-enriched, PR-enriched, LZV/PR-enriched, sweep, and the full set of SVs. Each sheet includes the GO ID, the name of the GO term, the GO domain, the function and the *P*-values and q-values.

**Supplemental Table 9.** Enrichment of insecticide resistance (IR) genes in structural variants (SVs) for each dataset. Insecticide resistance genes were significantly enriched in SVs in both the full set of SVs and the high impact set.

| Dataset | Number of real IR genes in dataset | Mean number of permuted IR genes | Number of instances where the number of IR genes in permutations ≥ real data | *P*-value |
|---|---|---|---|---|
| **High impact** | 91 | 48.129 | 0 / 10,000 | **0.0000** |
| **LZV-enriched** | 159 | 148.136 | 1,623 / 10,000 | 0.1623 |
| **Full Set** | 704 | 661.605 | 19 / 10,000 | **0.0019** |
| **Sweep** | 20 | 14.413 | 1,251 / 10,000 | 0.1251 |
| **High impact sweep** | 2 | 1.030 | 2,701 / 10,000 | 0.2701 |
| **PR-enriched** | 148 | 149.972 | 5,844 / 10,000 | 0.5844 |
| **LZV/PR-enriched** | 246 | 241.460 | 3,753 / 10,000 | 0.3753 |

**Supplemental Table 10.** Structural variants annotated by SnpEff as “5_prime_UTR_variant” and “3_prime_UTR_variant” and their corresponding insecticide resistance genes. Individual genotypes are also recorded, with ‘./’ indicating missing data.

**Supplemental Text:**

*Candidate insecticide resistance genes overlap genomic regions under positive selection*

With Illumina sequencing data from Haba et al. 2025, we identified signatures of recent selective sweeps in three *C. quinquefasciatus* populations using SweepFinder2 [1]. These populations, which included South Africa, Madagascar, and Cameroon/Gabon, were selected as proxies for our Zambian population. Selective sweeps were identified as regions where the CLR value exceeded the 99th percentile for the respective chromosome and population. We examined these regions for overlapping IR candidate genes and SVs, finding 13 total IR genes with SVs under putative positive selection. All genes highlighted here overlapped one or more SVs that were within a CLR peak and present in at least one Zambian individual.

In the South African population, we observed two IR genes overlapping CLR peaks (Supplemental Figure 33). On chromosome 1, two SVs within *tyramine beta hydroxylase* (LOC6053904) were also within a selective sweep region. Tyramine beta hydroxylase is involved in a pathway that is upregulated upon bendiocarb exposure in *Aedes aegypti* [2]. An additional peak on chromosome 2 overlapped four SVs within *retinol dehydrogenase 14* (LOC6054529). Retinol dehydrogenases have been identified as upregulated in SPI-exposed honeybee larvae [3].

In Madagascar, three IR gene SVs overlapped CLR peaks (Supplemental Figure 32)—although chromosome 1 lacked IR genes overlapping CLR peaks, peaks on chromosome 2 overlapped two uncharacterized genes (LOC6041432 and LOC6044529) with predicted roles in cuticle structure and oxidative stress response and peroxidase activity [4]. These genes contained one and seven SVs overlapping CLR peaks, respectively. On chromosome 3 in Madagascar, one IR gene overlapped a CLR peak. This gene was *xanthine dehydrogenase/oxidase* (LOC6047252), which contained two SVs overlapping the sweep. *Xanthine dehydrogenase/oxidase* encodes two enzymes involved in the response to oxidative stress [5], and duplication of an amplicon containing the *xanthine dehydrogenase* gene up to 250 copies has been observed in an organophosphate-resistant strain of *Cx. quinquefasciatus* [6].

In the Cameroon/Gabon population, eight total genes overlapped CLR peaks (Figure 4). On chromosome 1, *neuronal acetylcholine receptor subunit alpha-7* (LOC6051581) contained 17 SVs overlapping a peak. On chromosome 2, an *oxidase/peroxidase* gene (LOC6044529) contained eight SVs overlapping sweeps, and *probable cytochrome P450 12a5, mitochondrial* contained one SV overlapping a sweep. Finally, on chromosome 3, *cuticular protein 64Ac* (LOC6035808) overlapped three SVs within sweeps, *sulfide quinone oxidoreductase* (LOC6035804) overlapped two SVs within sweeps, *ionotropic receptor 75a* (LOC6047019) overlapped three SVs within sweeps, *Na channel protein 60E* (LOC6036601) overlapped 10 SVs within sweeps, and *peroxidasin* (LOC6047379) overlapped two SVs within sweeps. Neuronal acetylcholine receptors are well-known targets of neonicotinoid insecticides [7], and ionotropic receptors have also been shown to mediate IR [8,9]. Cytochrome P450s drive metabolic resistance by detoxifying insecticides [10]. Cuticle proteins have been implicated in IR in several species via increased cuticle thickness decreasing insecticide penetration [11,12]. In *Drosophila melanogaster*, *NaCP60E* modulates *para*-related pyrethroid resistance [13]. Oxidoreductase activity is enhanced in deltamethrin-resistant *Culex pipiens pallens* [14], and peroxidasin expression is upregulated in pyrethroid-resistant *Cimex lectularius* [15].

Within our sweep set of SVs, which was filtered for any SVs overlapping CLR peaks, we identified a total of 54 SVs that overlapped both IR genes and selective sweep regions. One of these SVs, a 728 bp deletion overlapping the 3' UTR of *cuticular protein 64Ac* (LOC6035808), was predicted by SnpEff to be high impact and result in a gene fusion between *cuticular protein 64Ac* and *neurobeachin-like protein 1* (LOC6035816). Although none of the remaining SVs that overlapped CLR outliers were classified as high impact by SnpEff, one overlapped the 5' UTR of *sulfide quinone oxidoreductase* (LOC6035804), which may impact the translation of its transcripts [16]. The remainder of the SVs overlapping these selective sweeps were contained within the introns of their respective IR genes.
